# Patterns of *Aedes aegypti* immature ecology and arboviral epidemic risks in peri-urban and intra-urban villages of Cocody-Bingerville, Côte d’Ivoire: insights from a dengue outbreak

**DOI:** 10.1101/2025.05.05.652274

**Authors:** Yasmine N. Biré, Julien Z. B. Zahouli, Jean D. K. Dibo, Pierre N. Coulibaly, Prince G. Manouana, Jacques F. Mavoungou, Fanny Hellhammer, Gaël D. Maganga, Luc S. Djogbenou, Ayola A. Adegnika, Steffen Borrmann, Stefanie C. Becker, Mahama Touré

**Affiliations:** Centre d’Entomologie Médicale et Vétérinaire, Université Alassane Ouattara, Bouaké, Côte d’Ivoire; Centre Suisse de Recherches Scientifiques en Côte d’Ivoire, Abidjan, Côte d’Ivoire; Centre de Recherches Médicales de Lambaréné, Lambaréné, Gabon; Institute for Tropical Medicine, University of Tübingen, Tübingen, Germany; Institut de Recherche en Ecologie Tropicale, Centre National de la Recherche Scientifique et Technologique, Libreville, Gabon; Institute for Parasitology University of Veterinary Medicine Hanover, Hanover, Germany; Research Center for Emerging Infection and Zoonoses, University of Veterinary Medicine Hanover, Hanover, Germany; German Center for Infection Research, Tübingen, Germany; Tropical Infectious Diseases Research Centre, University of Abomey-Calavi, Cotonou, Benin

## Abstract

**Background:** In Côte d’Ivoire, *Aedes* vector studies and arboviral outbreak responses have mostly focused on urbanized neighborhoods including intra-urban villages, but have often neglected peripheral villages. We assessed and compared *Aedes aegypti* population dynamics and dengue (DEN) and yellow fever (YF) epidemic risks among peri-urban and intra-urban villages during the 2023-2024 DEN outbreaks in Cocody-Bingerville, southern Côte d’Ivoire.

**Methods:** From August 2023 to July 2024, we sampled *Aedes* eggs, larvae and pupae in the domestic and peridomestic ecozones of three peri-urban and three intra-urban villages. We compared *Ae. aegypti* larval infestation, container productivity, *Stegomyia* indices (house index: HI, container index: CI, and Breteau index: BI) and pupal indices (pupae per house index: PHI, pupae per container index: PCI, and pupae per person index: PPI) between the study villages.

**Results:** *Aedes aegypti* widely dominated *Aedes* fauna in both peri-urban (98.1%) and intra-urban (99.8%) villages. Immatures were sampled from seven container types. However, the most productive containers were small containers, tires and medium containers which produced 81.8% of all the pupae in the peri-urban villages, and tires and small containers yielding 83.2% in the intra-urban villages. These key containers produced substantially more pupae in the domestic ecozones (70.9%) in the peri-urban villages, but equitably between the domestic (48.8%) and peridomestic (51.2%) ecozones in the intra-urban villages. In each study village, they yielded around 90% and 80% of pupae during the short dry and long rainy seasons, respectively. CI, HI and BI were comparable between the peri-urban (29.9%, 35.9% and 41.4) and intra-urban (36.7%, 48.0% and 56.2) villages, and surpassed the World Health Organization (WHO) DEN and YF epidemic thresholds. PCI (1.26 *vs.* 3.38 pupae/container) and PHI (1.75 *vs.* 5.18 pupae/house) were lower in the peri-urban than in the intra-urban villages, while PPI (0.54 *vs.* 1.25 pupae/person) was similar between the villages. Unlike temperature and relative humidity, all indices were strongly associated with rainfall in all the study villages.

**Conclusion:** In Cocody-Bingerville, all sampled peri-urban and intra-urban villages exhibited high densities of *Ae. aegypti* immatures and habitats. Although larval habitats were diverse, a limited number of container types accounted for over 80% of pupal production. *Stegomyia* indices remained consistently elevated, exceeding WHO epidemic thresholds for DEN and YF across all study villages, potentially contributing to ongoing DEN outbreaks. Therefore, vector surveillance and outbreak responses should be extended to peri-urban villages, as they could act as arbovirus re-introduction sources. Community-based interventions targeting the identified key containers could help control the outbreaks. This study provides a baseline for strategically reducing human exposure to arboviruses in the study villages.

## Introduction

*Aedes* mosquito-borne arboviruses, including dengue virus (DENV), yellow fever virus (YFV), chikungunya virus (CHIKV) and Zika virus (ZIKV), are posing significant threats to the public and global health worldwide [1,2]. Arboviral diseases are posing a significant threat to over 831 million people (approx. 70% of population) on the continent [3,4]. In 2023 alone, 171,991 suspected cases of dengue (DEN), including 70,223 cases and 753 deaths were reported from 15 African countries, including Côte d’Ivoire [5]. In Africa, arboviruses are mostly transmitted to humans by *Aedes aegypti* and *Aedes albopictus* species [3]. The spread and geographical expansion of *Ae. aegypti* and *Ae. albopictus* and arboviruses are mainly driven by urbanization and climate change [3,4].

Rapid and uncontrolled urbanization in Africa provide millions of water-holding containers and solid or plastic waste (e.g., water storage receptacles, tires, cans, etc.) which can act as *Aedes* ovipositing and breeding sites, and humans that are blood-feeding sources for the females [6,7]. Breeding sites are filled with water manually by people or naturally by rainfalls making them suitable for ovipositing and larval development. Other meteorological variables, such as high temperature and high relative humidity (RH), can accelerate larval development and increase adult survivorship [8,9]. Thus, fast urbanization and climate warming are expected to increase *Aedes* adult density and biting rate and exacerbate the arboviral transmission and burden in Africa [3,7]. However, no licensed vaccines are still available for many arboviruses (except for yellow fever (YF) vaccine) in most African countries [3,10]. Therefore, the surveillance of *Aedes* vectors is crucial for the preparedness for and the response, prevention and control of arboviral epidemics. Côte d’Ivoire is one of most important emerging and re-emerging foci of DEN and YF in Africa [11,12]. Four DENV serotypes (i.e., DENV1-4) and YFV are present there. In the recent years (2017-2024), the country has faced multiple DEN and YF outbreaks, with the majority (80-90%) of cases reported in the region of Abidjan (7 million inhabitants) [13–15]. For instance, DEN outbreaks caused 321 infected cases and 27 deaths in 2023 [16], and 4,450 confirmed and 2 fatal DEN cases in 2024 [17]. These numbers likely represent only a fraction of the actual cases, as many may remain undiagnosed or unreported. Nonetheless, over 80-90% of DEN and YF cases reported in Abidjan are recorded in the health district of Cocody-Bingerville (∼900,000 inhabitants). Outbreaks mostly occurred during rainy seasons in Cocody-Bingerville. Data highlight the ongoing and expanding threat of the diseases in both urban and peri-urban areas.

In Cocody-Bingerville, urbanization is rapidly expanding into rural areas, thus drawing many intra-urban and peri-urban villages. Despite this, as arboviral cases are predominantly reported in the urban hospitals, leading most studies on local *Aedes* vectors and arboviral risk assessment to focus primarily on the urban areas and intra-urban villages, often overlooking the peri-urban villages [6]. As a result, outbreak responses (e.g., removals of larval breeding sites, insecticide space spraying against *Aedes* larvae and adults, and local community awareness) are mostly implemented in the urban areas and intra-urban villages [11,12]. Though, numerous peri-urban villages are closely interconnected with and strongly influenced by these highly urbanized areas. Indeed, significant movement of people and items (e.g., tires and cans) occurs between the urban areas, the intra-urban villages and the peri-urban villages. However, the peri-urban villages often faced limited access to adequate water supply and waste management services, which are more readily available urbanized areas. This lack of infrastructure can lead to the storage of water and the accumulation of unmanaged containers that can create breeding sites for the aquatic stages of *Aedes* mosquitoes [14, 18], Such conditions can increase the density of *Ae. aegypti* adult populations and biting rates, and the risk of DEN and YF epidemics in the peri-urban areas, similar to those observed in urban and intra-urban villages of Cocody-Bingerville [19]. Despite these risks, few studies have examined *Aedes* vectors and arboviral epidemic risks in the peri-urban villages.

The present study aimed to assess and compare the patterns of *Ae. aegypti* larval infestation and container productivity and associated DEN and YF epidemic risks among peri-urban and intra-urban villages during the 2023-2024 DEN outbreaks in Cocody-Bingerville. To do so, we monitored *Aedes* immature forms (eggs, larvae and pupae) and evaluated the level of the epidemic risks of DEN and YF using both qualitative (*Stegomyia* indices) and quantitative (container productivity) approaches [20,21]. The findings provided crucial insights into entomological factors influencing DENV and FYV transmission and offer valuable data to support the development of targeted control strategies in Cocody-Bingerville, a key arboviral hotspot in Côte d’Ivoire.

## Methods

### Study area

The study was carried out in peri-urban villages and intra-urban villages within the health district of Cocody-Bingerville (population ∼ 897,000 inhabitants) in southeastern Côte d’Ivoire (Fig 1). This district is a part of the lagoon region administrated by the economic capital city of Abidjan (population ∼ 7 million inhabitants), Rapid and ongoing urban expansion characterizes the area, resulting in a heterogeneous landscape that includes numerous peri-urban and intra-urban settlements, from which the present study sites were selected. Cocody-Bingerville represents currently the major DEN and YF hotspot of the country, and harbor high densities of *Aedes* mosquitoes and abundant breeding sites [6,22,23]. Local arboviral incidences and *Ae. aegypti* population typically peak during the rainy seasons [6,22,23].

**Fig. 1.**
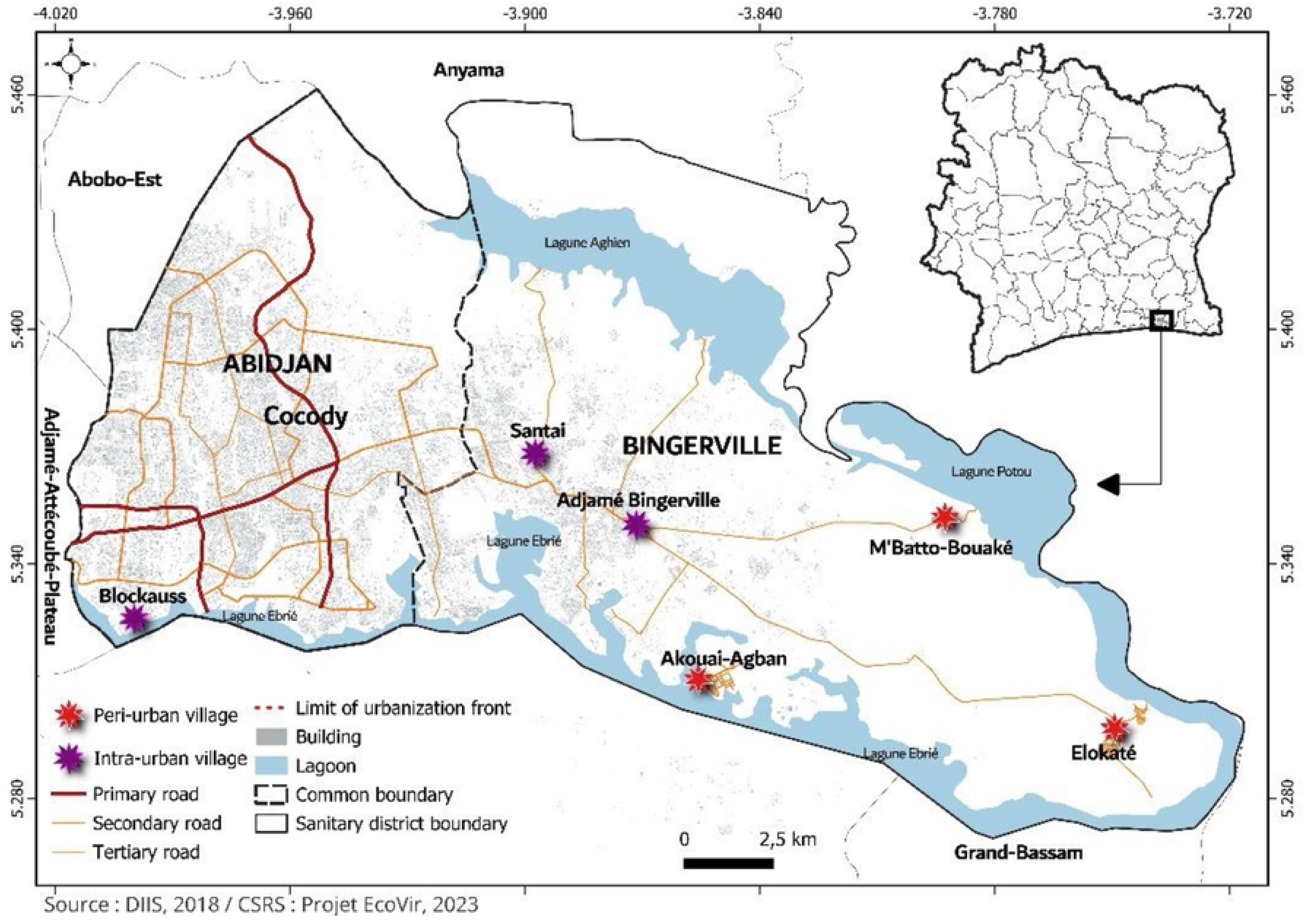
Location of the study sites located in the Heath district of Cocody-Bingerville, southeastern Côte d’Ivoire from August 2023 to July 2024. The study sites were peri-urban villages (A) and intra-urban villages of Cocody-Bingerville. The peri-urban villages comprise Akouai-Agban, M’batto-Bouaké and Elokaté characterized by secondary paved roads and mixture of green spaces, agricultural areas and spaced houses, and are located 5-15 km from the borders of the urban areas of Cocody-Bingerville. The intra-urban villages include Adjamé-Bingerville, Blockhauss and Santé that are all urban neighborhoods with numerous major and secondary paved roads and dense blocks of flats and located within the highly urbanized parts of the municipalities of Cocody and Bingerville.

Cocody-Bingerville is in a coastal area with a tropical climate characterized by high temperatures and high RH throughout the year. The climate is marked by four rainfall-based seasons: long dry season (LDS) from December to March, long rainy season (LRS) from April to July, short dry season (SDS) from August to September, and short rainy season (SRS) from October and November. Average annual precipitation ranges from 1,200 to 2,400 mm. The annual temperature is approximately 26.5 °C. The annual RH ranges between 78 and 90%.

### Study design

This study was conducted in three peri-urban villages and three intra-urban villages, totaling six study sites with roughly equal size. The peri-urban villages were represented by the villages of Akouai-Agban (5° 18’ 39” N, 3° 51’ 20” W), Elokaté (5° 17’ 38” N, 3° 44’ 56” W) and M’batto-Bouaké (5° 19’ 0” N, 3° 48’ 2” W) (Figure 1). All sites were located within a 15 km radius from the border of the urban health district of Cocody-Bingerville. The selected sites were characterized by a mixture of residential, commercial and agricultural land-use, with infrastructure limited to unpaved or secondary paved roads. The peri-urban villages (6,300 inhabitants) are less densely populated than the intra-urban areas (9,200 inhabitants) [24]. The intra-urban villages included Adjamé-Bingerville (5° 20’ 45” N, 3° 52’ 20” W), Blockhauss 5° 19’ 24” N, 4° 0’ 7” W) and Santé (5° 21’ 56” N, 3° 53’ 55” W) (Figure 1). These intra-urban villages were located within the urban municipalities of Abidjan and Bingerville, and were characterized by advanced infrastructure, including residential and commercial buildings, primary and paved roads and presence of trade, industrial and administrative services. Each study site was subdivided into two ecological zones: domestic and peridomestic ecozones. Domestic ecozones were defined as spaces characterized by the presence of human-inhabited houses and associated structures. Peridomestic ecozones referred to areas extending up to 50 m surrounding the domestic ecozones. They included public or private spaces, such as roadsides, garages, schools, green spaces, agricultural fields and bushes.

Collections were done quarterly from August 2023 to July 2024. Cross-sectional surveys were conducted during SDS, LRS, SRS and LDS at each study site, resulting in a total of 24 collection events (12 in peri-urban and 12 in intra-urban villages).

### Egg collections

*Aedes* mosquito eggs were collected using standard WHO ovitrap method [25,26]. A total of 100 ovitraps (50 in domestic ecozone and 50 in peridomestic ecozone) was placed at each study site during each survey. Therefore, 2,400 ovitraps (1,200 in per-urban villages and 1,200 in intra-urban villages) deployed. The ovitraps were cut-out metal and back-painted boxes with a volume of 400 cm^3^. Each ovitrap was garnished with hardboard paddles, rough on one side which served as an egg-laying substrate. The ovitraps were filled out at ¾ volume with a mixture of distilled water and rainwater (ratio: 1:1) [25] and were left in the field for a one-week during each survey. All ovitrap contents (paddles, mosquito larvae or pupae and water) were collected and transferred separately into different plastic cups labelled (household code, ecozone type, study site and collection date).

### Larval and pupal collections

We searched for and sampled *Aedes* mosquito larvae and pupae among water-holding containers. Collections were done in 100 households (i.e., in domestic and peridomestic ecozones) randomly selected per study site and per survey. All pupae were collected. The larval and pupal samples were stored separately in plastic cups labelled (household code, ecozone type, study site and collection date). All inspected breeding sites were characterized and classified into seven categories according to the WHO guideline [27], as follows: 1) Large containers: drums and barrels (i.e., water volume > 50 liters); 2) Medium containers (i.e., buckets, larges pots, and small barrels: volume between 10-49 liters); 3) Small containers (i.e., all container types: Water volume <10 liters); 4) Tires (i.e., discarded bicycle, motorcycle, car, or other motor vehicle tires); 5) Troughs (i.e., any type of container of any material used to water animals); 6) Flowerpots (i.e., flower containers); and 7) Others (i.e., brick holes, shoes, tarpaulins, wooden boxes, mortar, sheet metal, leaf armpits, snail shells, puddles and tree holes).

### Laboratory procedures

All collected mosquito samples (eggs, larvae and pupae) were transported in cool boxes to the field laboratory. Each sample was put in plastic cup labelled with the sampling site, ecozone and date. The paddles were dried for seven days and immersed separately in distilled water for egg hatching. All mosquito larvae were fed with Tetra-Min Baby Fish Food. Egg-derived and field-collected larvae and pupae were raised to adults under ambient conditions. Emerged adults were identified morphologically to species and sex under a binocular magnifying glass using available taxonomic keys [25,28]. Data was recorded in a designed entomological database.

### Climate data

Data on the climatic conditions, including rainfall, temperature and RH, were obtained from the Société d’Exploitation et de Développement Aéroportuaire, Aéronautique et Météorologique (SODEXAM) of Côte d’Ivoire (www.sodexam.com).

### Statistical analysis

Data were analyzed using R Studio version 4.4.2. A significance level of 5% was set. The container infestation rate (IR) was expressed as the percentage of containers with at least one *Ae. aegypti* larva or pupa (numerator) among the wet containers (denominator). The proportion of positive breeding site types (PP) was estimated as the percentage of each *Ae. aegypti*-positive container type (numerator) among the total *Ae. aegypti*-positive containers (denominator). Chi-square test (χ^2^) was used to compare FP and PP between the study areas.

*Aedes aegypti* oviposition indices such as ovitrap positivity index (OPI: percentage of *Ae. aegypti*-positive ovitraps among total number of ovitraps), mean egg count per ovitrap (MEO: mean number of *Ae. aegypti* eggs per number of ovitraps) and egg density index (EDI: mean number of *Ae. aegypti* eggs per *Ae. aegypti-*positive ovitraps) [29] were each compared between study areas and across seasons using generalized linear mixed model (GLMM) with Poisson family. DEN and YF outbreak risks were determined using *Ae. aegypti* larval infestation levels based on *Stegomyia* indices that included container index (CI: percentage of *Ae. aegypti*-positive containers among water-holding containers), house index (HI: percentage of houses with at least one *Ae. aegypti*-positive container) and Breteau index (BI: number of *Ae. aegypti*-positive containers per 100 houses). HI, CI and BI were compared across the study areas using one-way analysis of variance (ANOVA). Moreover, HI, CI and BI values were compared with the WHO-established DEN and YF epidemic thresholds. DEN epidemic risk is high if CI > 3% or HI > 4% and BI > 5 [30]. YF epidemic risk is low if IC < 3%, HI < 4% or BI < 5; moderate if 3% ≤ CI ≤ 20%, 4% ≤ HI ≤ 35% or 5 ≤ BI ≤ 50; and high if CI > 20%, HI > 35% or BI > 50 [31]. *Aedes aegypti* container productivity was estimated through pupal indices: pupae per house index (PHI: number of pupae divided by the number of houses), pupae per container index (PCI: number of pupae divided by the number of containers) and pupae per person index (PPI: number of pupae divided by the number of persons living in the houses). Additionally, as the *Ae. aegypti* pupal mortality was very low, human biting index (HBI) was calculated as the number of females emerged from pupae divided by the number of persons dwelling in the inspected houses [32]. PHI, PCI, PPI and HBI were compared between the study areas using ANOVA.

We tested the relationship between oviposition indices and *Stegomyia* indices using the Pearson test. Additionally, correlations between pupal indices, *Ae. aegypti* indices and the local climate variables (i.e., rainfall, temperature and RH) using the Sperman’s test, as the data were not normally distributed (Shapiro-Wilk test was significant, p < 0.05).

### Ethics statement

Before sample collection, the study protocol received ethical approval from the national ethics committee of Côte d’Ivoire (N/Ref: 071-23/MSHPCMU/CNESVS-km). Additionally, permissions were obtained for the General Directorate of Health (GDH) of the Ministry of Health (MoH) of Côte d’Ivoire, the administrative and health authorities of Cocody-Bingerville and the local community leaders of the study areas. Mosquito samples and data were collected with the authorization of the residents and/or owners. This study did not involve endangered or protected species.

## Results

### Climate indicators

Rainfall, temperature and RH were high and varied across seasons in the peri-urban and intra-urban villages (Table 1). During the surveys, the respective averages for these climate variables were 147.00 mm of rainfall, 27.90 °C temperature and 80.20% RH. Rainfall, temperature and RH varied significantly over seasons (rainfall: F = - 04.14, df = 3, p < 0.0001; temperature: F = 105,09, df = 3, p < 0.0001; and RH: F = 133,40, df = 3, p < 0.0001). Rainfall was highest during LRS (329.90 mm) and lowest during SDS (17.05 mm). The temperature peaked during LDS (30.92 °C) and was lowest during LRS (26.50 °C). RH was highest during LRS (84.50%) and lowest during SDS (75.19%).

**Table 1.**
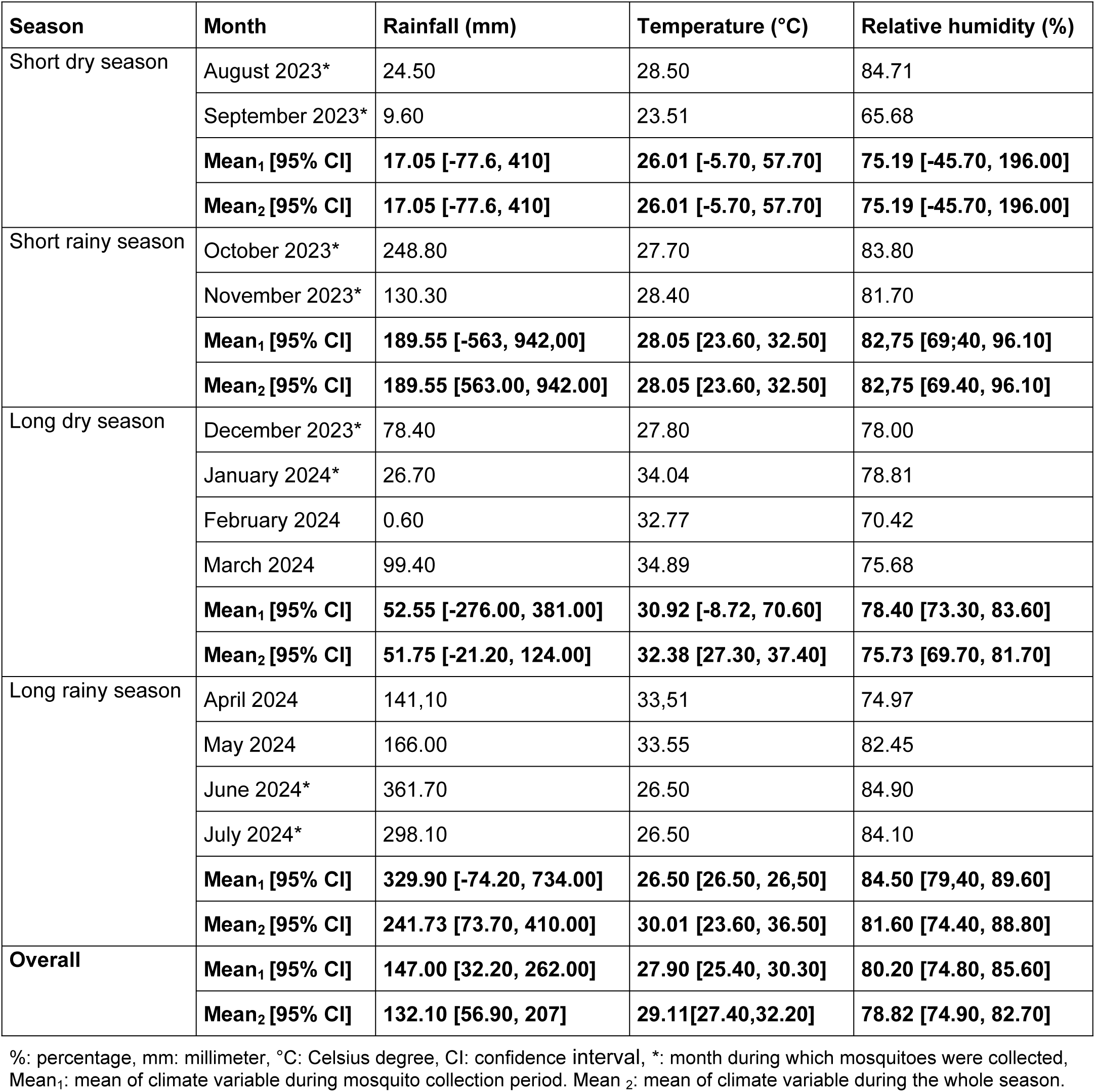
Climate indicators recorded seasonally in the heath district of Cocody-Bingerville, Côte Ivoire, from August 2023 to July 2024.

### Mosquito species composition

*Aedes aegypti* widely dominated *Aedes* genus in both peri-urban villages (98.05%, 13,157/13,419) and intra-urban villages (99.78%, 29,168/29,232). Taken together, *Ae. aegypti* proportion was 2.2-time lower in the peri-urban villages (31.09%, 13,157/42,325) compared to the intra-urban villages (68.91%, 29,168/42,325) (Table 2). Moreover, *Ae. aegypti* number was significantly lower in the peri-urban villages than in the intra-urban villages (χ^2^ = 81.570, df = 1, p < 0.0001). In the peri-urban villages, five additional *Aedes* species were found, among which *Aedes vittatus* (1.39%, 187/13,419) showed a proportion above 1% (Table 2). Key non-*Aedes* mosquito taxa of public health importance, including *Culex quinquefasciatus* and *Eretmapodites* species *(Eretmapodites chrysogaster* and *Eretmapodites quinquevittatus)* vectors of arboviruses were mostly collected in the peri-urban villages, while *Anopheles gambiae* malaria vector was mainly encountered in the intra-urban villages (S1 Table).

**Table 2.** Species composition of *Aedes* mosquito adults emerged from eggs, larvae and pupae sampled in the peri-urban and intra-urban villages of Cocody-Bingerville, southeastern Côte d’Ivoire, from August 2023 to July 2024.

| Village | Species | Eggs |  |  |  | Larvae |  |  |  | Pupae |  |  |  | Total |  |  |  |
| --- | --- | --- | --- | --- | --- | --- | --- | --- | --- | --- | --- | --- | --- | --- | --- | --- | --- |
|  |  | Female | Male | Total | % | Female | Male | Total | % | Female | Male | Total | % | Female | Male | Total | % |
| Peri-urban | *Aedes aegypti* | 2768 | 2445 | 5213 | 99.58 | 2787 | 3061 | 5848 | 96.90 | 1032 | 1064 | 2096 | 97.53 | 6587 | 6570 | 13157 | 98.05 |
|  | *Aedes dendrophilus* | 5 | 16 | 21 | 0.40 | 0 | 0 | 0 | 0 | 0 | 0 | 0 | 0 | 5 | 16 | 21 | 0.16 |
|  | *Aedes fraseri* | 0 | 0 | 0 | 0 | 5 | 1 | 6 | 0.10 | 5 | 8 | 13 | 0.60 | 10 | 9 | 19 | 0.14 |
|  | *Aedes lillii* | 0 | 0 | 0 | 0 | 8 | 12 | 20 | 0.33 | 0 | 0 | 0 | 0 | 8 | 12 | 20 | 0.15 |
|  | *Aedes luteocephalus* | 0 | 0 | 0 | 0 | 2 | 13 | 15 | 0.25 | 0 | 0 | 0 | 0 | 2 | 13 | 15 | 0.11 |
|  | *Aedes vittatus* | 1 | 0 | 1 | 0.02 | 70 | 76 | 146 | 2.42 | 24 | 16 | 40 | 1.86 | 95 | 92 | 187 | 1.39 |
|  | **Total** | **2774** | **2461** | **5235** | **100** | **2872** | **3163** | **6035** | **100** | **1061** | **1088** | **2149** | **100** | **6707** | **6712** | **13419** | **100** |
| Intra-urban | *Aedes aegypti* | 4410 | 4862 | 9272 | 99.99 | 6935 | 6730 | 13665 | 99.80 | 2882 | 3349 | 6231 | 99.43 | 14227 | 14941 | 29168 | 99.78 |
|  | *Aedes fraseri* | 0 | 0 | 0 | 0 | 21 | 6 | 27 | 0.20 | 1 | 2 | 3 | 0.05 | 22 | 8 | 30 | 0.10 |
|  | *Aedes vittatus* | 0 | 1 | 1 | 0.01 | 0 | 0 | 0 | 0 | 8 | 25 | 33 | 0.53 | 8 | 26 | 34 | 0.12 |
|  | **Total** | **4410** | **4863** | **9273** | **100** | **6956** | **6736** | **13692** | **100** | **2891** | **3376** | **6267** | **100** | **14257** | **14975** | **29232** | **100** |

?: percentage.

### *Aedes aegypti* population dynamics

#### Population abundance

*Aedes aegypti* abundance varied across the ecozones and seasons in the peri-urban and intra-urban villages. *Aedes aegypti* proportions were significantly higher in the domestic ecozones (61.94%, 8,149/13,157) compared with the peridomestic ecozones (38.06% (5,008/13,157) in the peri-urban villages (χ^2^ = 1498.8, df = 1, p < 0.0001). However, the proportions were statistically comparable between the domestic ecozones (49.95%, 14,568/29,168) and the peridomestic ecozones (50.05%, 14,600/29,168) in the intra-urban villages (χ^2^ = 0.0659, df = 1, p = 0.7974) (S2 Table). The highest proportions in the peri-urban and intra-urban villages were recorded during LRS, with respective peaks of 45.95% (6,045/13,157) and 39.19% (11430/29,168) (Fig 2). Similarly, the lowest proportions in the peri-urban villages (10.75%, 1,414/13,157) and the intra-urban villages (12.70%, 3,704/29,168) were observed during SDS. The proportions in LDS were significantly higher compared with SDS in the peri-urban (χ^2^ = 956.47, df = 2, p < 0.0001) and intra-urban villages (χ^2^ = 3774.3, df = 2, p < 0.0001). Surprisingly, the proportions in LDS were significantly higher than that recorded in SDS and SRS in both peri-urban villages (SDS: χ^2^ = 953.02, df = 1, p < 0.0001; SRS: χ^2^ = 217.34, df = 1, p < 0.0001) and intra-urban villages (SDS: χ^2^ = 2618.6, df = 1, p < 0.0001; SRS: χ^2^ = 1156.6, df = 1, p < 0.0001). Similar patterns were observed in the domestic and peridomestic ecozones in the peri-urban and intra-urban villages, with pronounced magnitude in the domestic ecozone of both villages (Fig 2). During any season, the proportions were significantly higher in the domestic ecozones compared with the peridomestic ecozones in the peri-urban villages (all p < 0.05), but statistical similar between the two ecozones in the intra-urban villages (all p > 0.05) (Fig 3). In the domestic ecozones, *Ae. aegypti* proportions and breeding sites in the peri-urban villages were higher than that in the intra-urban villages during all the seasons (Fig 3). Conversely, *Ae. aegypti* proportions in the peridomestic ecozones were lower in the peri-urban villages compared with the intra-urban villages during any season (Fig 3A).

**Fig 2.**
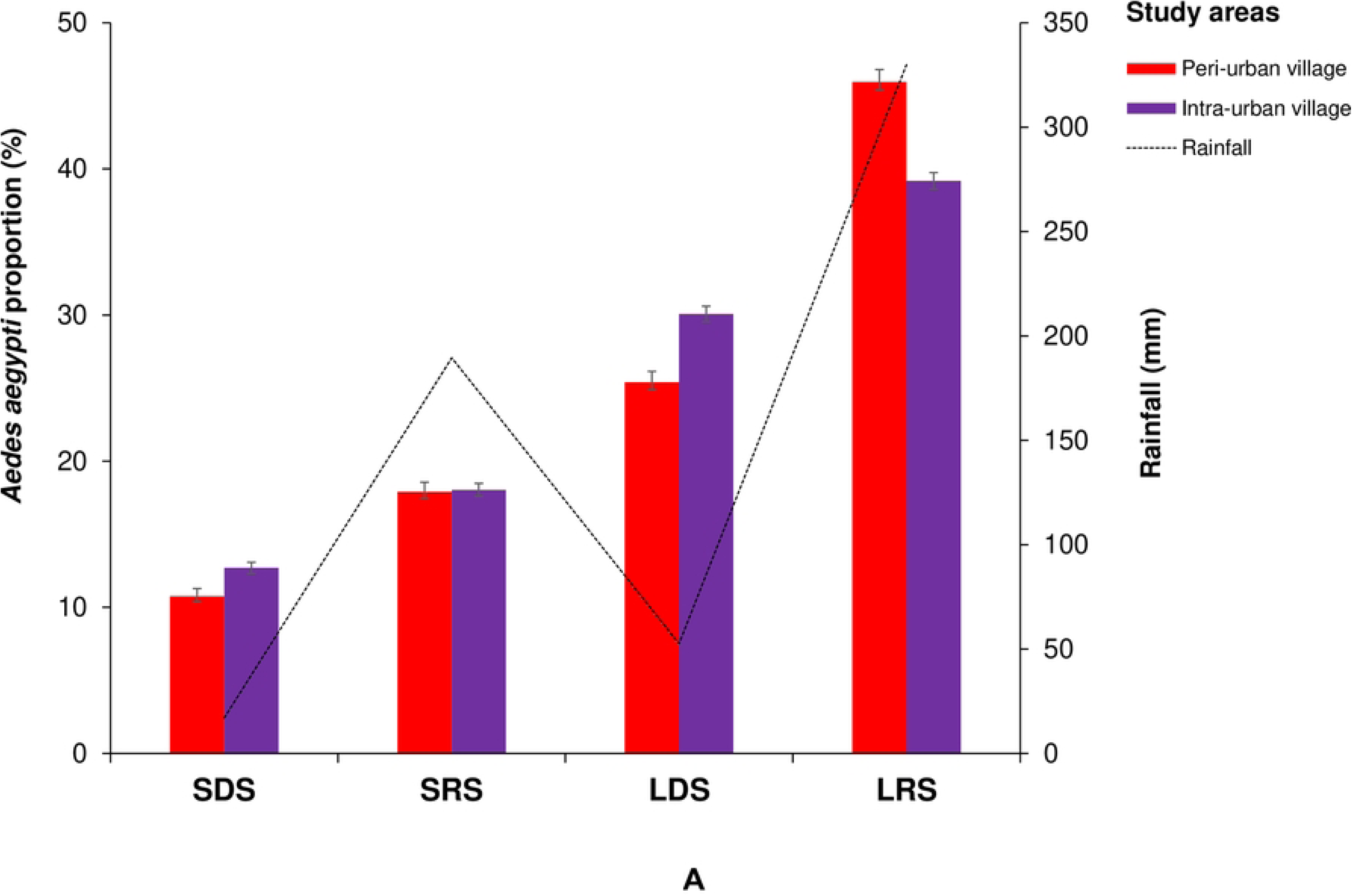
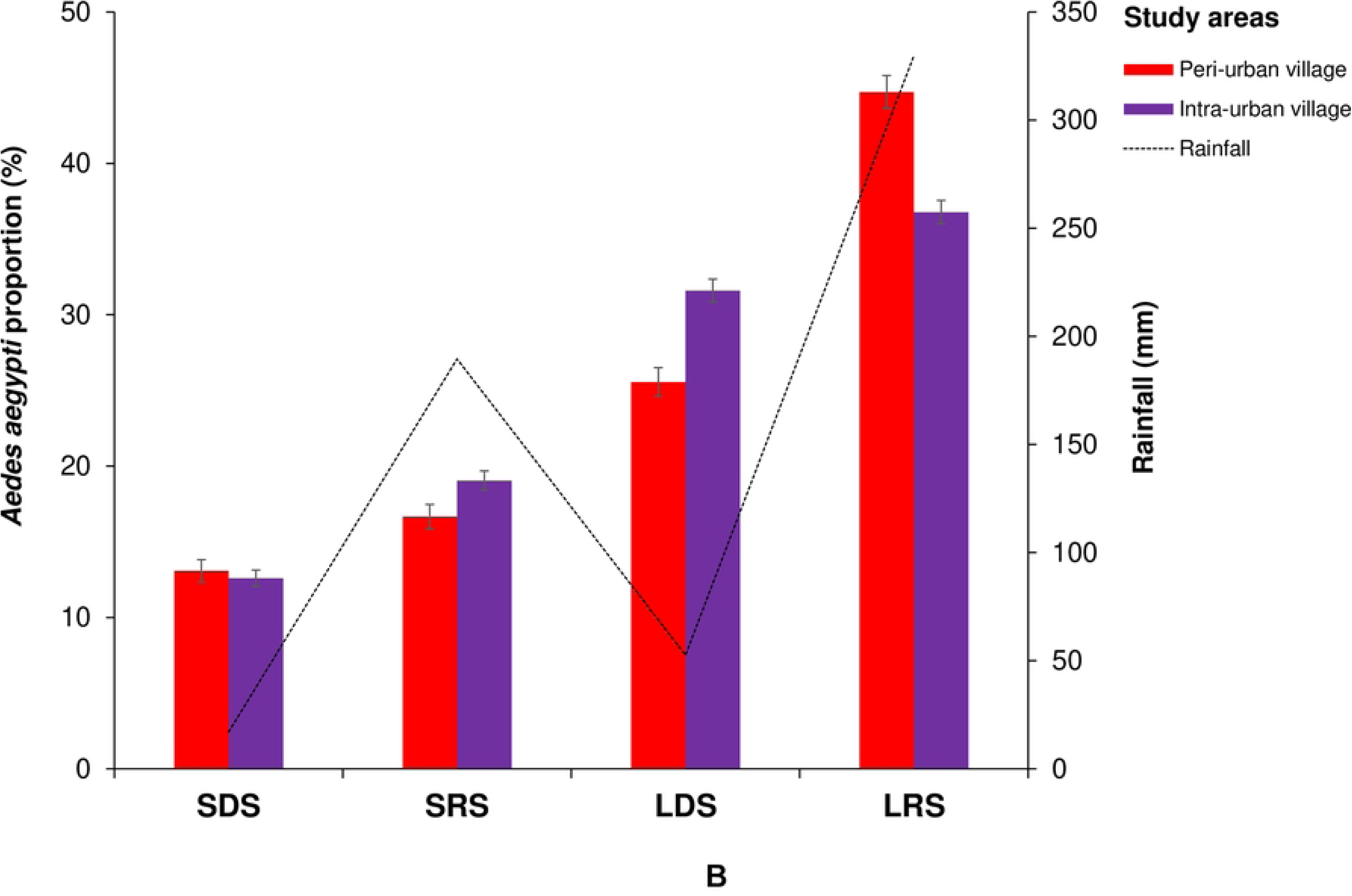
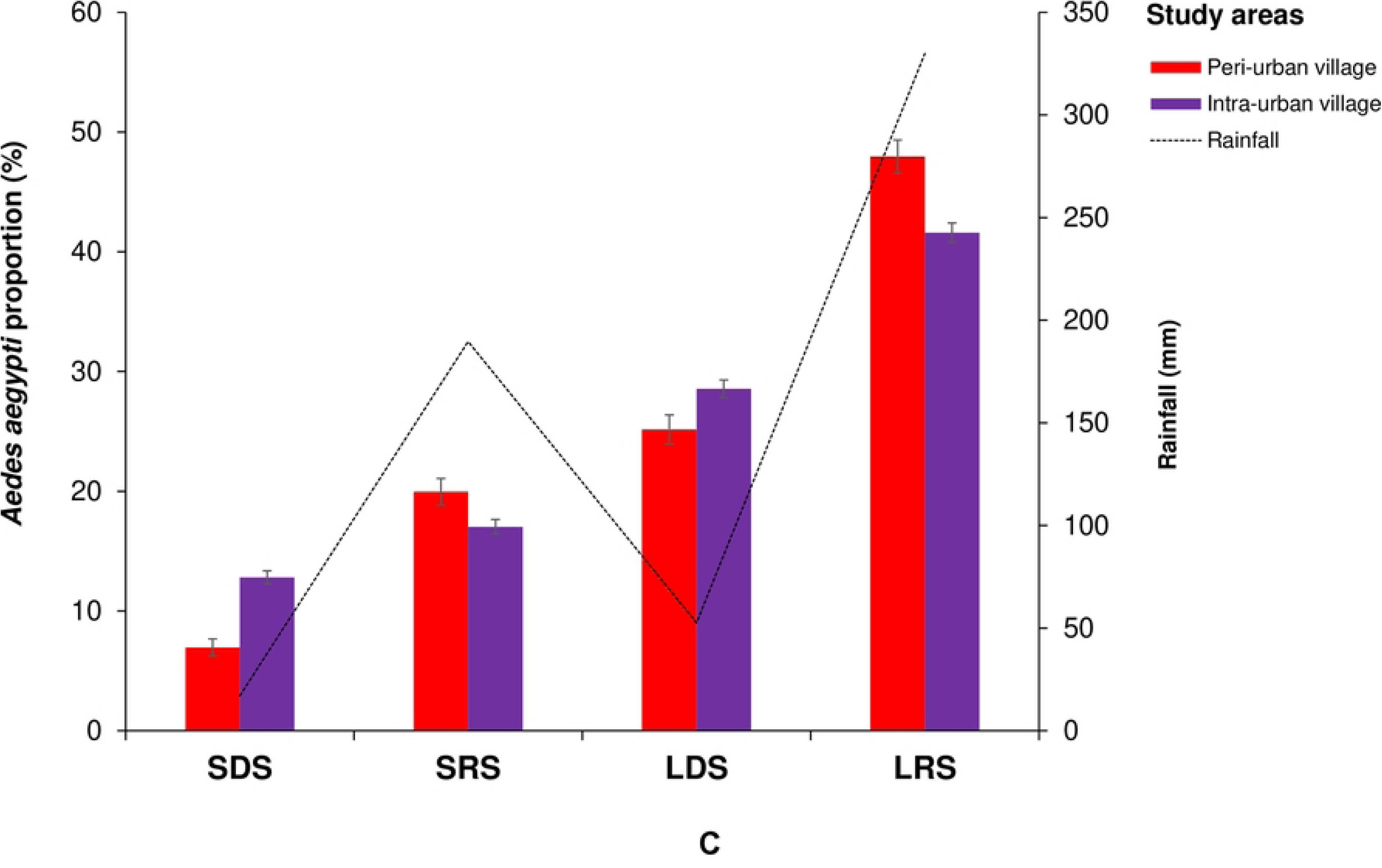
Seasonal variations in the proportions of *Aedes aegypti* adults emerged from eggs, larvae and pupae sampled in the peri-urban and intra-urban villages of Cocody-Bingerville, southeastern Côte d’Ivoire, from August 2023 to July 2024. A: Overall, B: Domestic ecozone, C: Peridomestic ecozone. LDS: long dry season, LRS: long rainy season, SDS: short dry season, SRS: short rainy season. Error bars show confidence intervals (95% CI).

**Fig 3.**
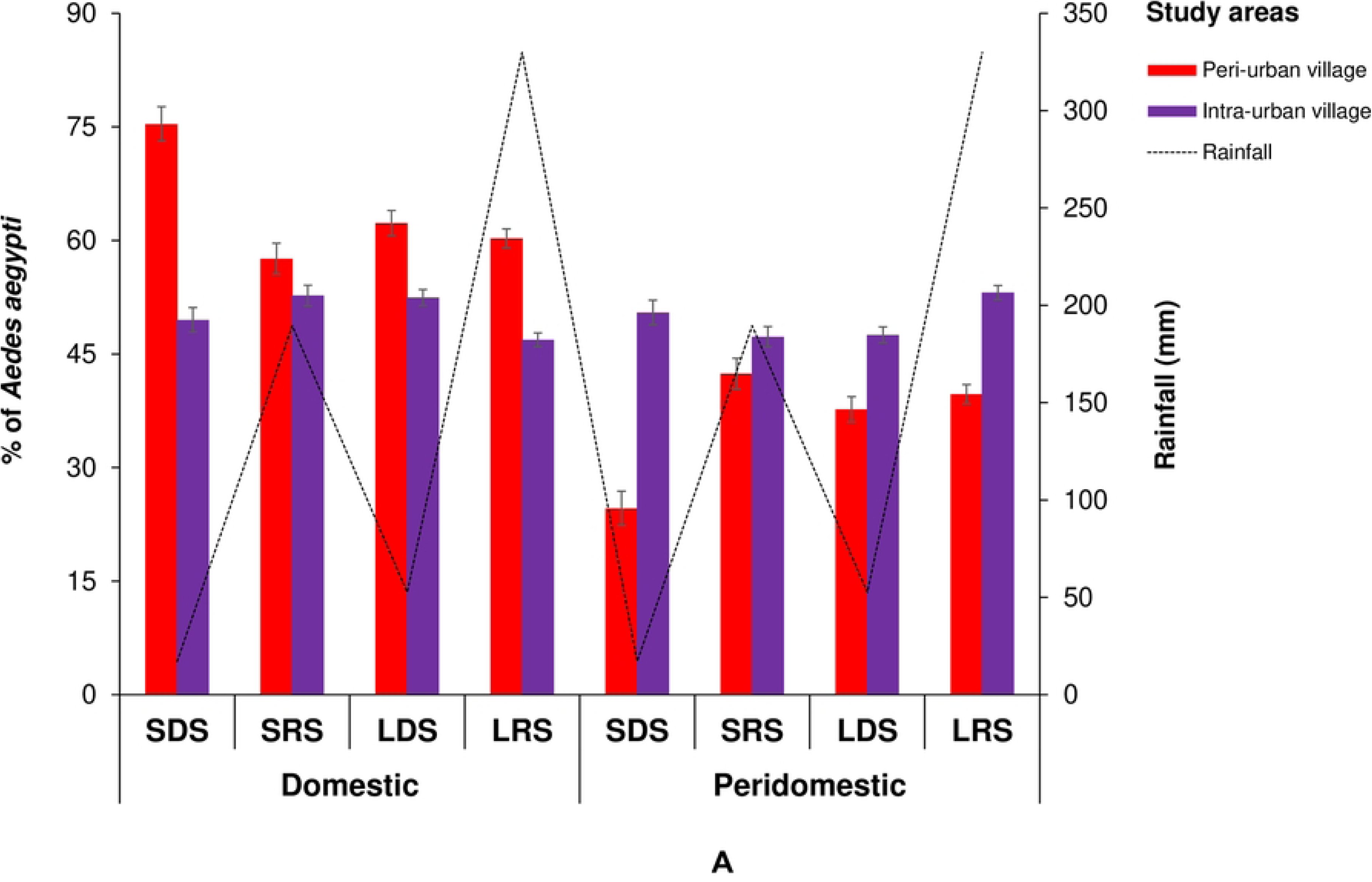
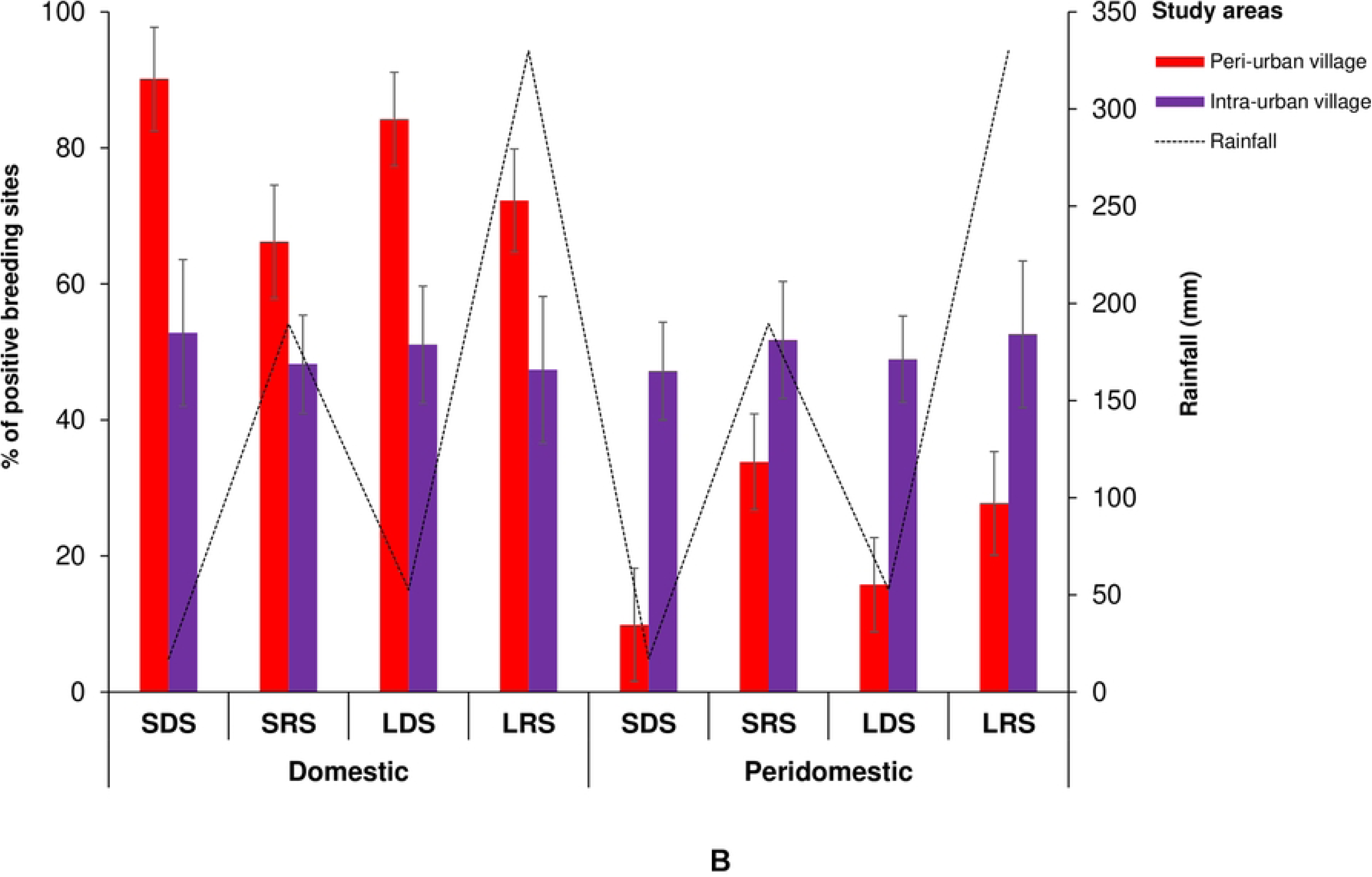
Geographical variations in *Aedes aegypti* proportions and breeding site infestation in the domestic and peridomestic ecozones in the peri-urban and intra-urban villages of Cocody-Bingerville, southeastern Côte d’Ivoire, from August 2023 to July 2024. A: *Aedes aegypti* proportion, B: Breeding site proportion. LDS: long dry season, LRS: long rainy season, SDS: short dry season, SRS: short rainy season. Error bars show confidence intervals (95% CI).

#### Larval infestation

The *Ae. aegypti*-infestation rates of containers were significantly lower in the peri-urban villages (IR = 29.96%, 497/1,659) compared with the intra-urban villages (IR = 36.71%, 674/1,836) (χ^2^ = 17.534, df = 1, p < 0.0001) (S3 Table). All the seven types of containers were found infested with *Ae. aegypti* immatures in both study villages. Container-specific positivity rates varied by types of villages (Fig 4). In the peri-urban villages, the most abundant *Ae. aegypti*-positive containers were small containers (PP = 34.41%, 171/497), tires (PP = 30.38%, 151/497) and medium containers (PP = 14.29%,71/497). In the intra-urban villages, tires dominated with PP of 63.65%, 429/674), followed by small containers (PP = 18.55%, 125/674) (Fig 5). Pooled together, *Ae. aegypti*-positive containers were significantly more abundant in the peri-urban villages (PP = 42.44%, 497/1,171) than in the intra-urban villages (PP = 57.56%, 675/1,171) (χ^2^ = 52.905, df = 1, p < 0.0001).

**Fig 4.**
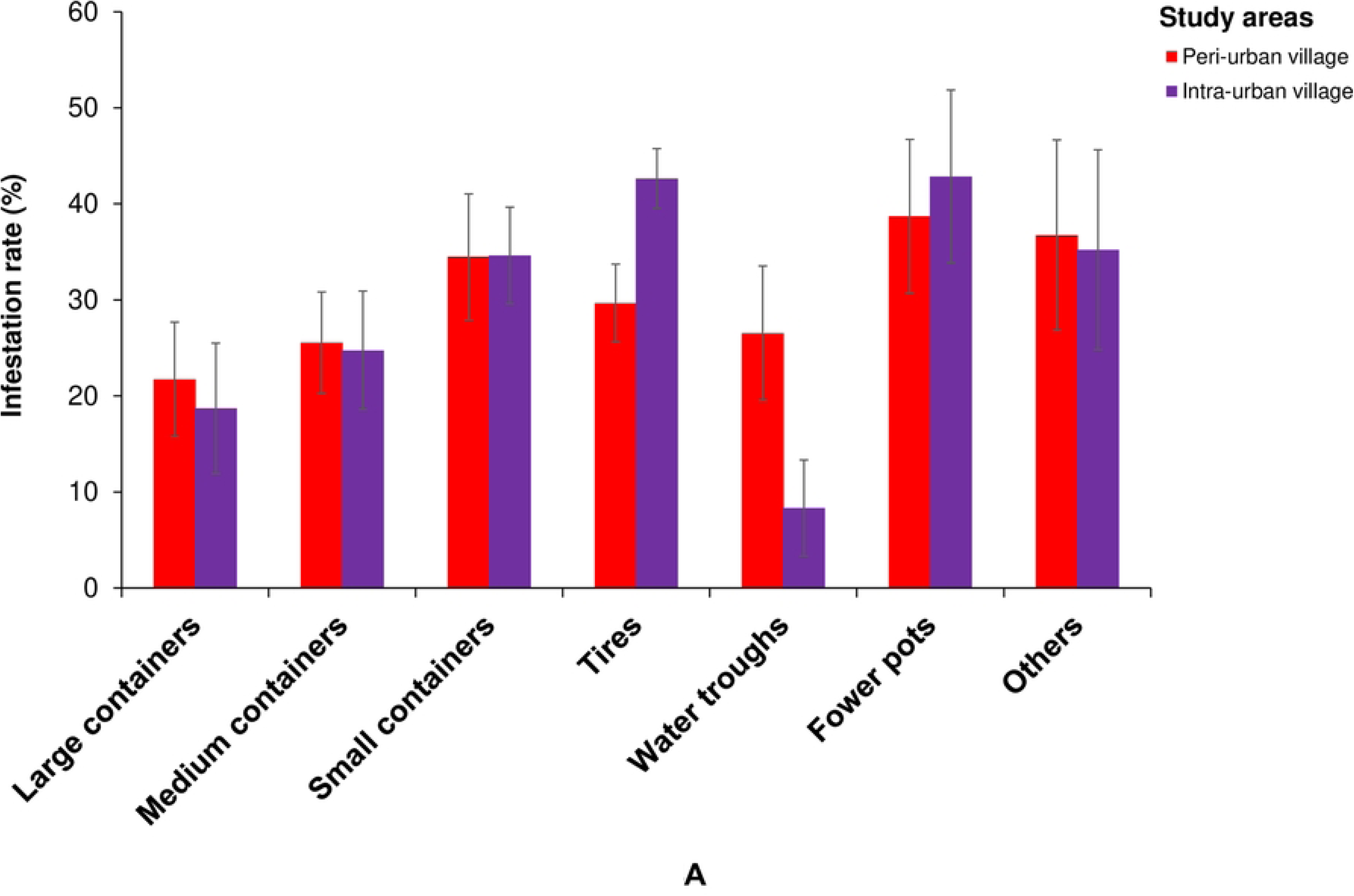
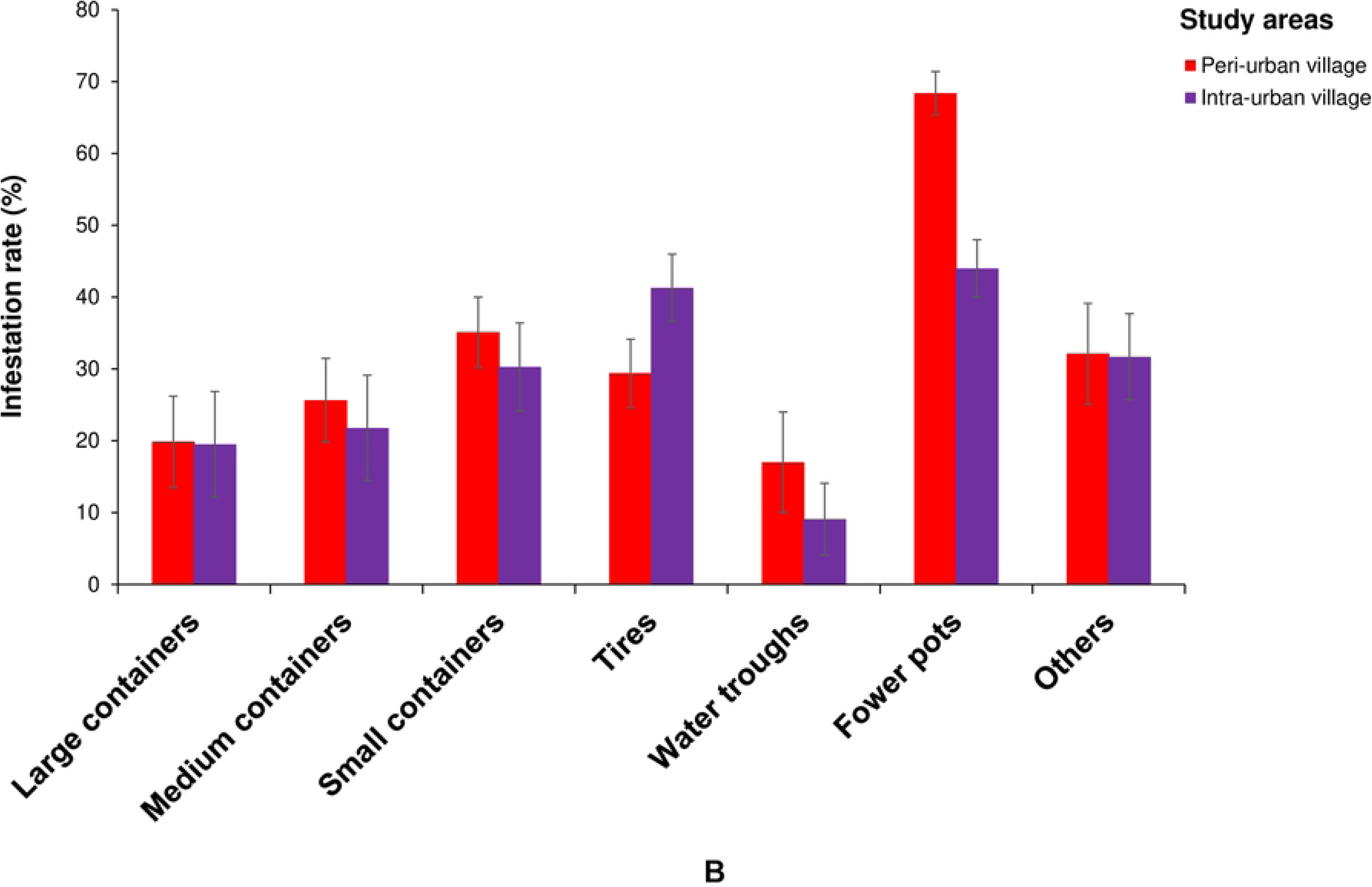
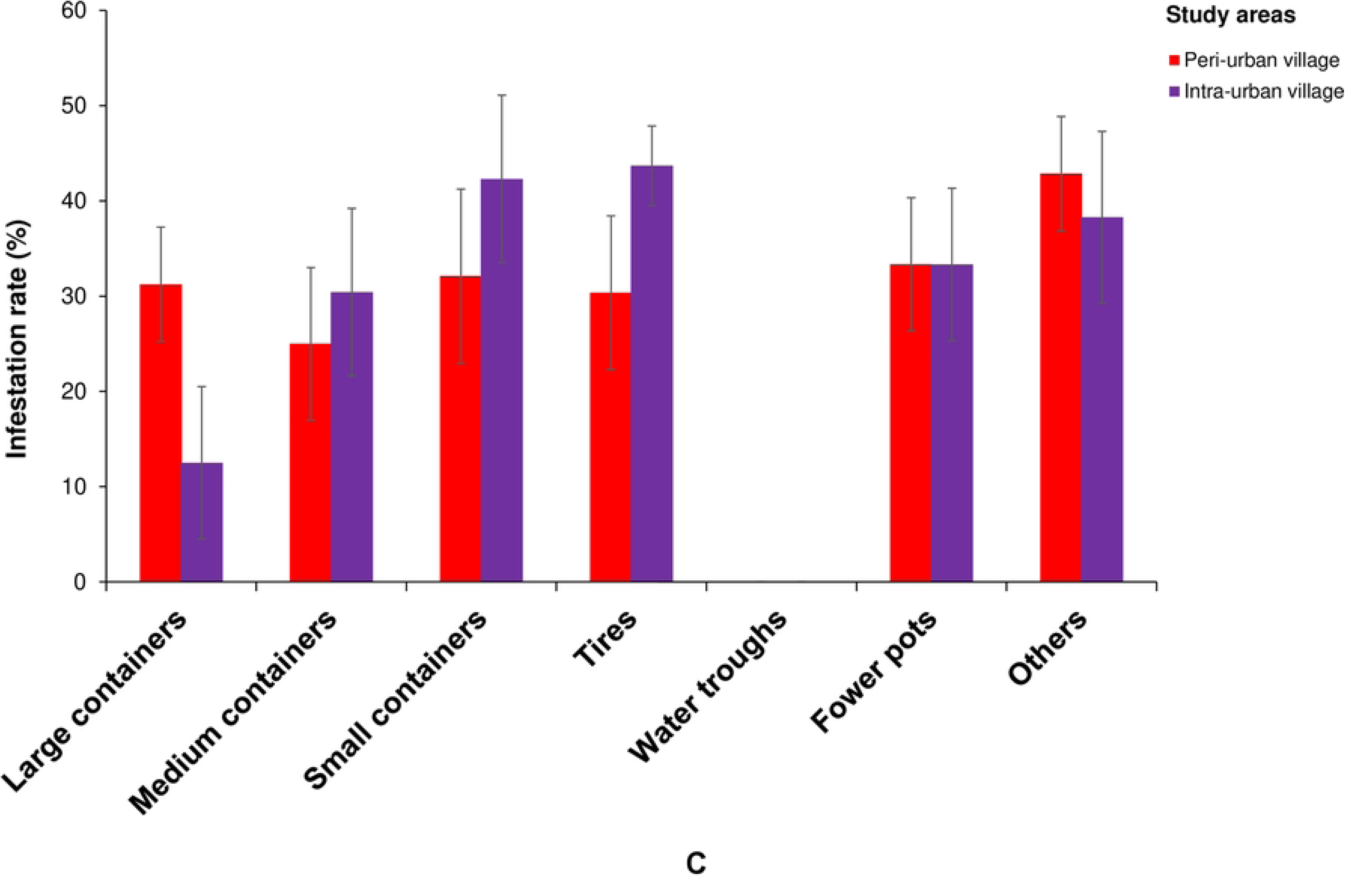
Infestation of *Aedes aegypti* breeding sites in the peri-urban and intra-urban villages of Cocody-Bingerville, southeastern Côte d’Ivoire, from August 2023 to July 2024. A: Overall, B: Domestic ecozone, C: Peridomestic ecozone. Error bars indicate confidence intervals (95% CI). Others includes breeding containers made with brick holes, shoes, tarpaulins, wooden boxes, mortar, sheet metal, leaf armpits snail shells, underground puddles and tree holes.

**Fig 5.**
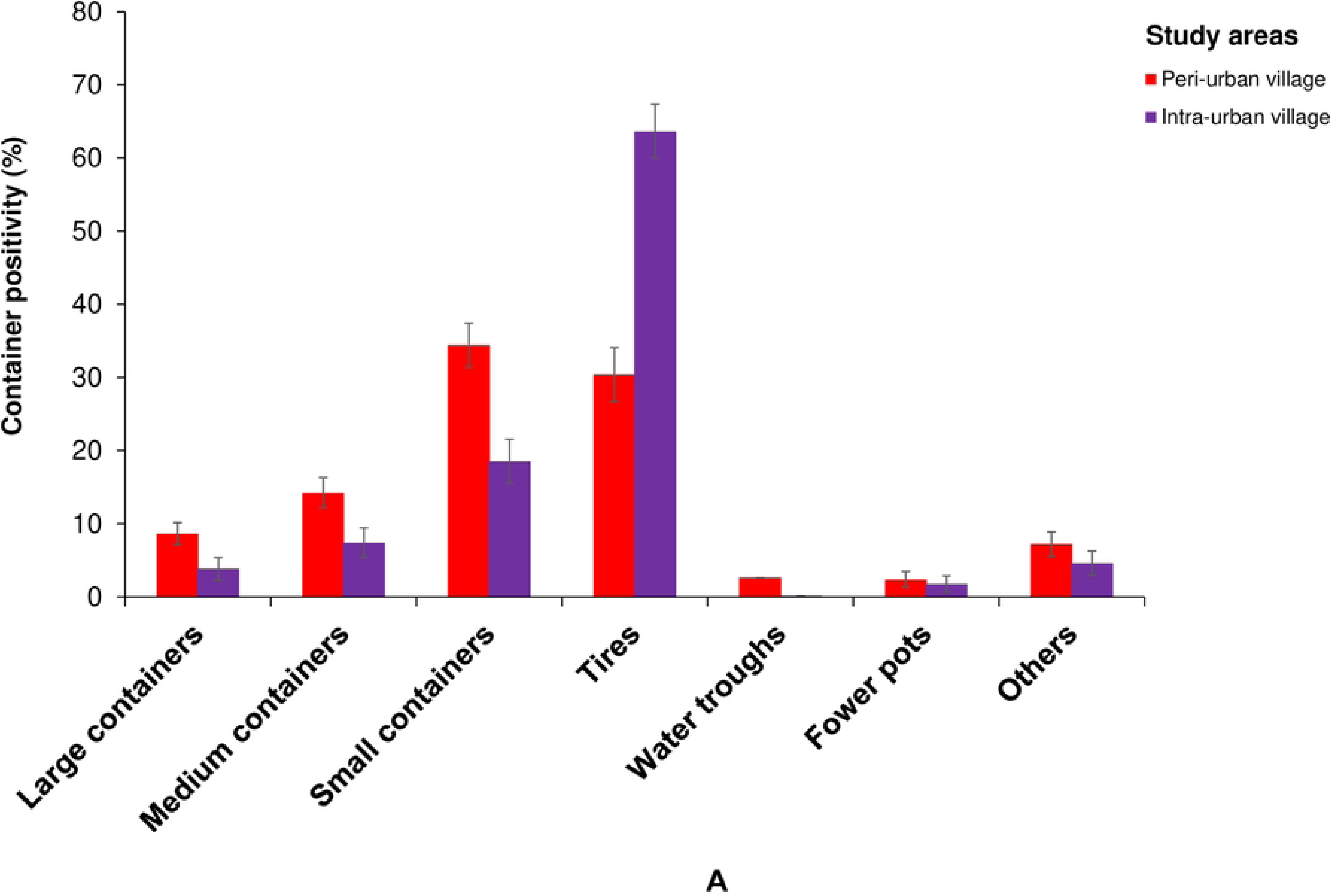
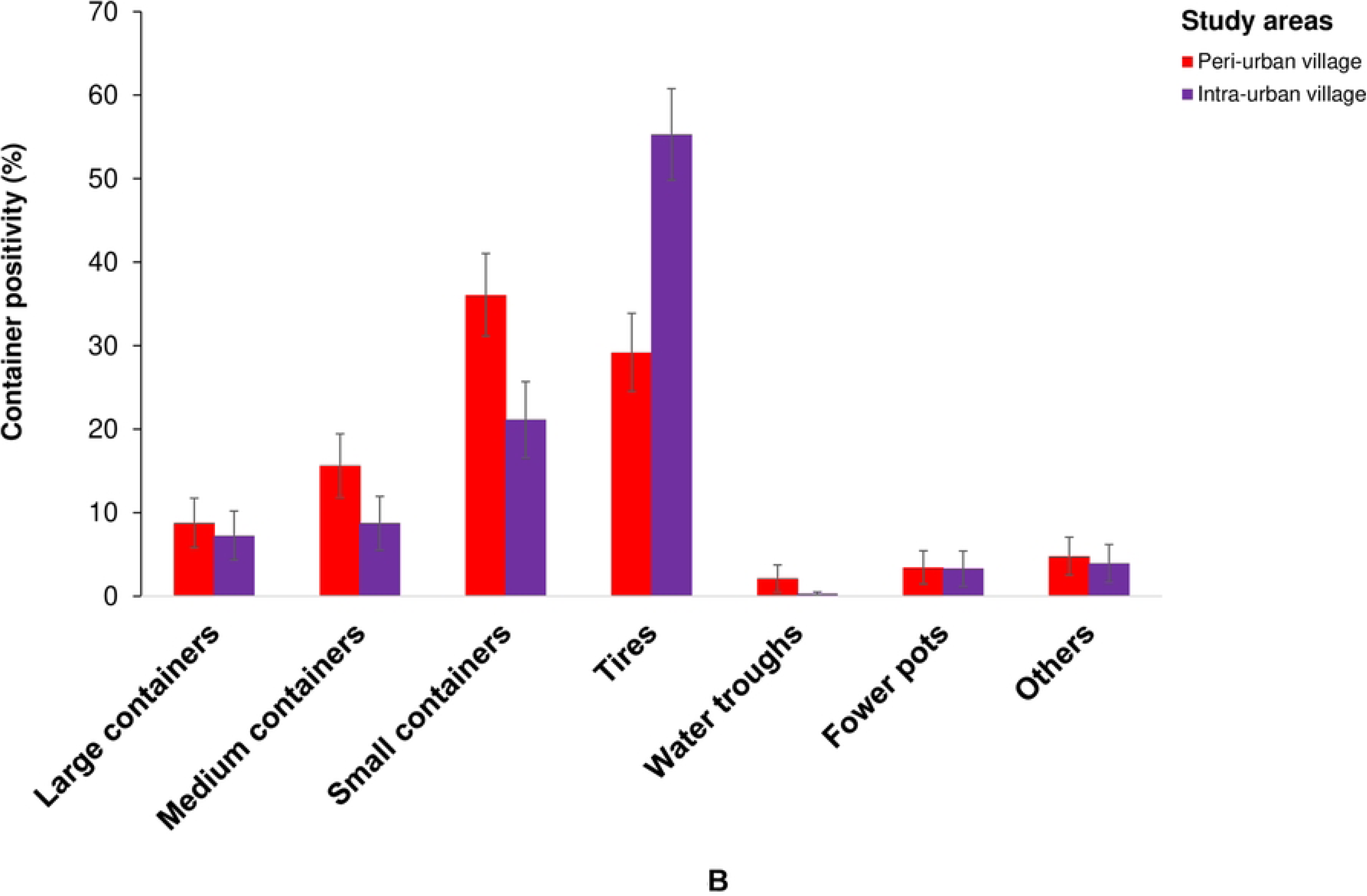
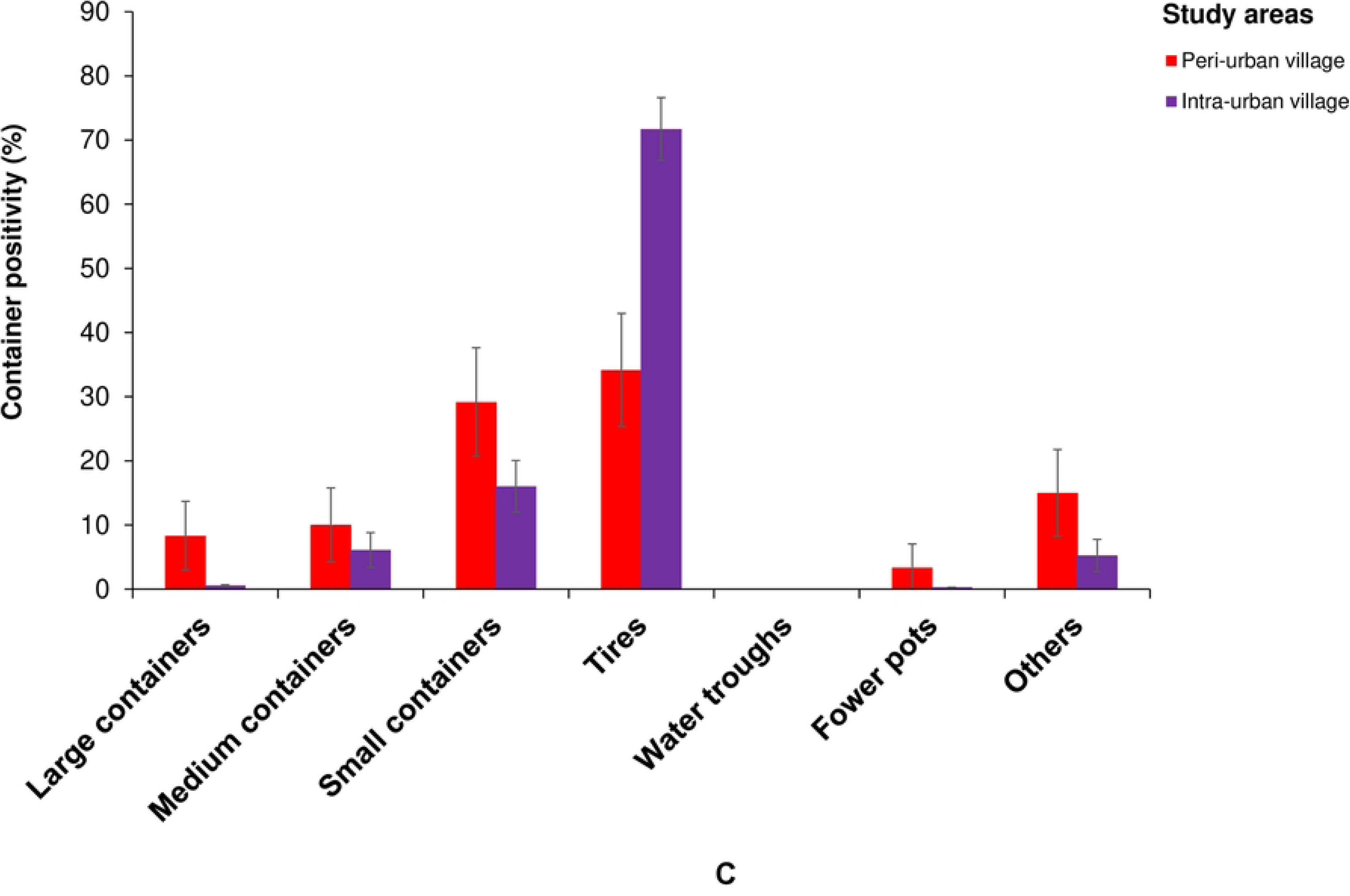
Proportion of *Aedes aegypti* breeding sites in the peri-urban and intra-urban villages of Cocody-Bingerville, southeastern Côte d’Ivoire, from August 2023 to July 2024. A: Overall, B: Domestic ecozone, C: Peridomestic ecozone. Error bars indicate confidence intervals (95% CI). Others includes breeding containers made with brick holes, shoes, tarpaulins, wooden boxes, mortar, sheet metal, leaf armpits snail shells, underground puddles and tree holes.

The proportions of *Ae. aegypti*-infested containers differed across the ecozones and the seasonal variability in all the study villages. The positive containers’ proportions were 3-time and significantly higher in the domestic ecozones (75.55%, 377/497) compared to the peridomestic ecozones (24.14%, 120/497) in the peri-urban villages (χ^2^ = 263.73, df = 1, p < 0.0001). *Ae. aegypti*-positive container proportions were nearly equal between the domestic ecozones (49.11%, 331/674) and the peridomestic ecozones (50.89% (343/674) in the intra-urban villages (χ^2^ = 0.3591, df = 1, p = 0.549). Seasonal patterns showed more pronounced variability in the peri-urban villages, with positive containers’ proportions peaking during SDS in the domestic ecozones and during SRS in the peridomestic ecozones. In contrast, the peri-urban villages showed relatively similar proportions of positive containers in both domestic and peri-domestic ecozones across seasons. Compared with the intra-urban villages, the peri-urban villages yielded consistently more positive containers in the domestic ecozones, but a lower number in the peridomestic ecozones across all seasons (Fig 3B).

#### Container productivity

Containers produced 3-fold lower number of *Ae. aegypti* pupae in the peri-urban villages (2,096 pupae) than in the intra-urban villages (6,231 pupae) (χ^2^ = 4104.7, df = 1, p < 0.0001). In the peri-urban villages, the most productive containers were small containers (31.11%, 652/2,096), followed by tires (30.53%, 640/2,096) and medium containers (20.13%, 422/2,096), providing together 81.77% (1,714/2,096) of the total pupae (Fig 6A). In the intra-urban villages, the key containers were tires (64.45%, 4,001/6,231) and small containers (18.75%, 1,164/6,231) that yielded 83.20% (5,165/6,231) of the total pupae (Fig 6A). Container productivity during LDS was significantly higher compared with that found during SDS in the peri-urban villages (χ^2^ = 160.56, df = 1, p < 0.0001) and the intra-urban villages (χ^2^ = 896.33, df = 1, p < 0.0001), and that recorded during SRS in both peri-urban (χ^2^ = 13.165, df = 1, p < 0.0001) and intra-urban (χ^2^ = 272.49, df = 1, p < 0.0001) villages (Table 3). Similar patterns were observed in the domestic and peridomestic ecozones in the peri-urban and intra-urban villages, but with pronounced magnitude in the domestic ecozones

**Fig 6.**
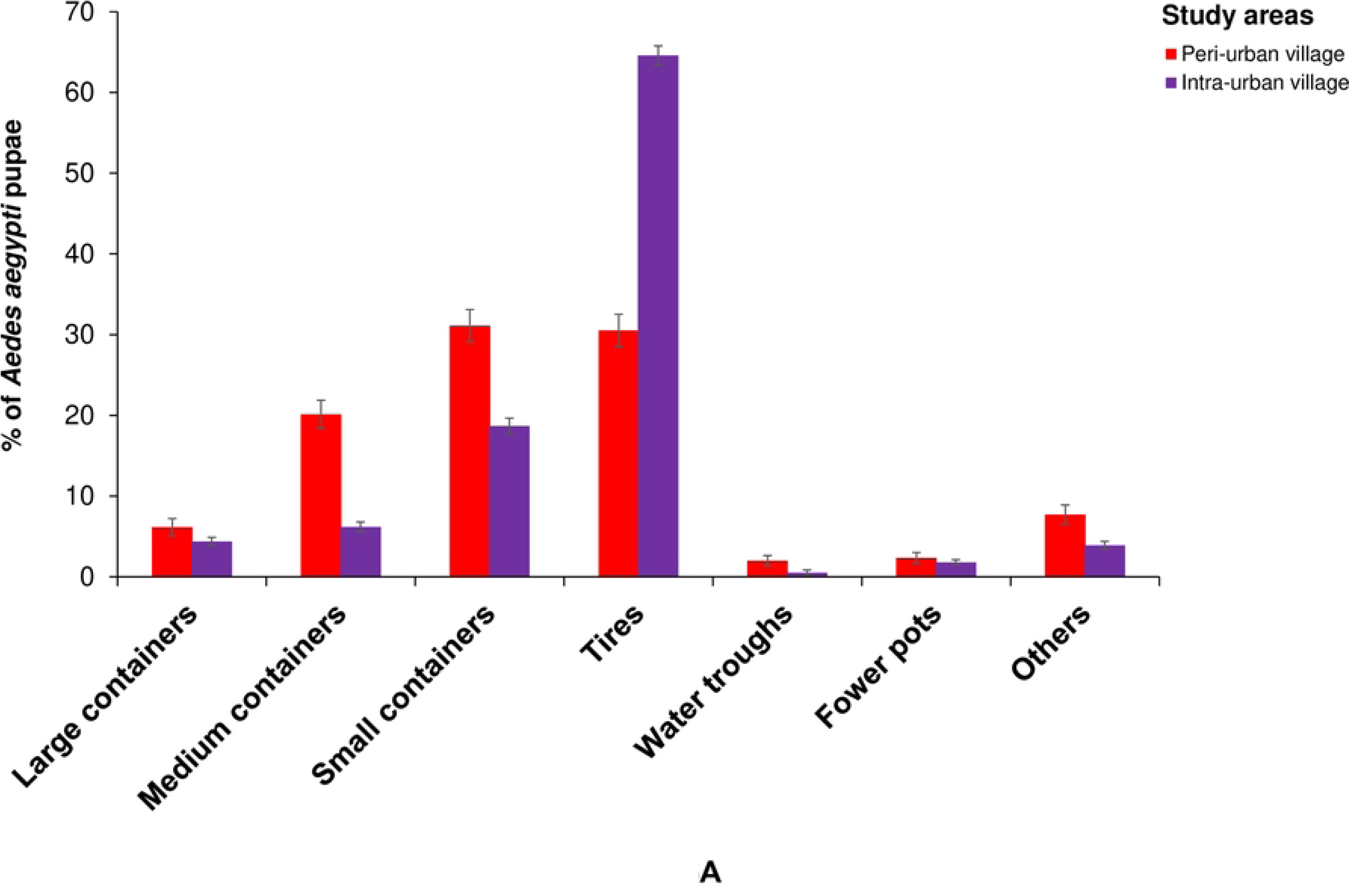
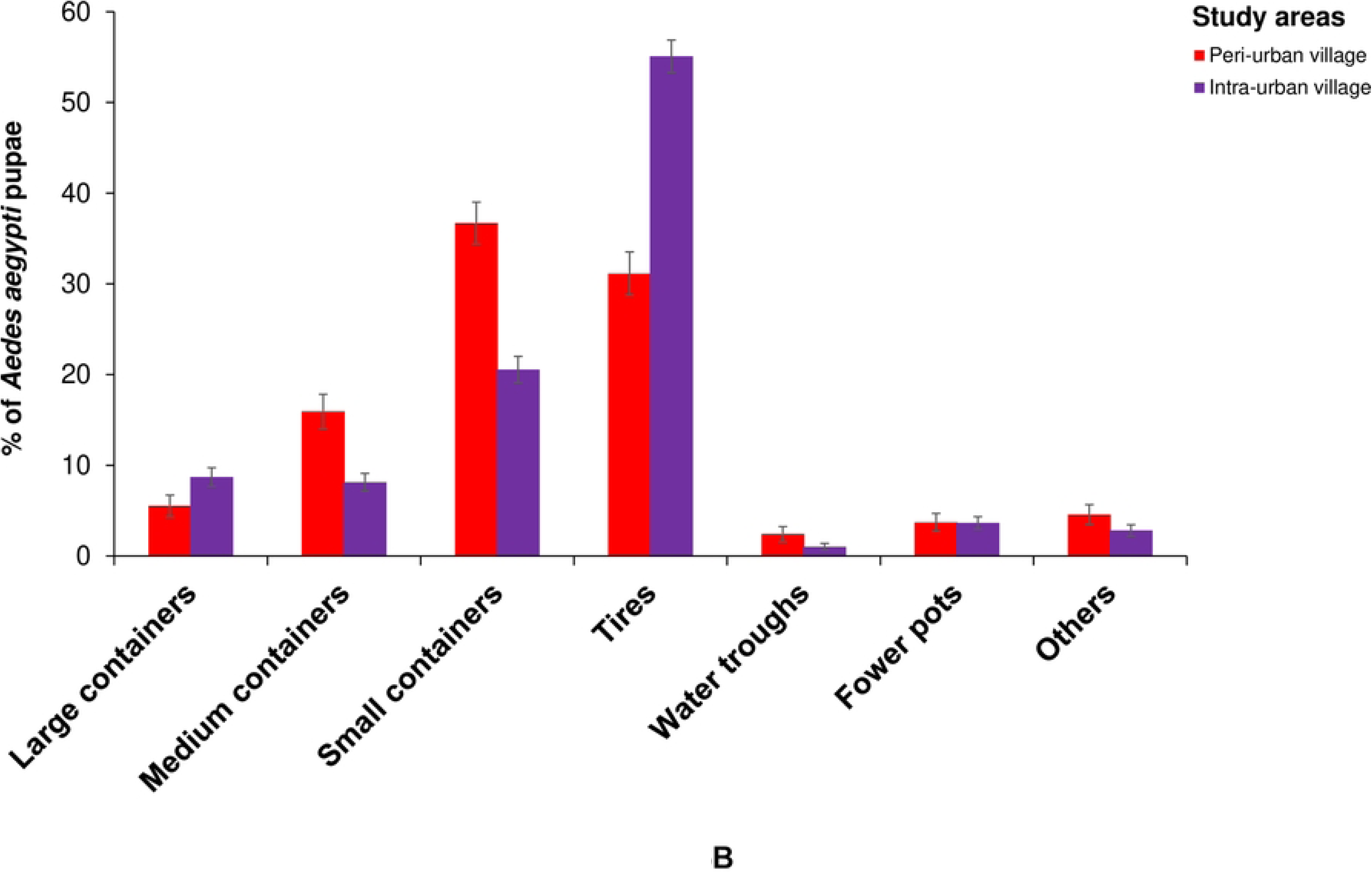
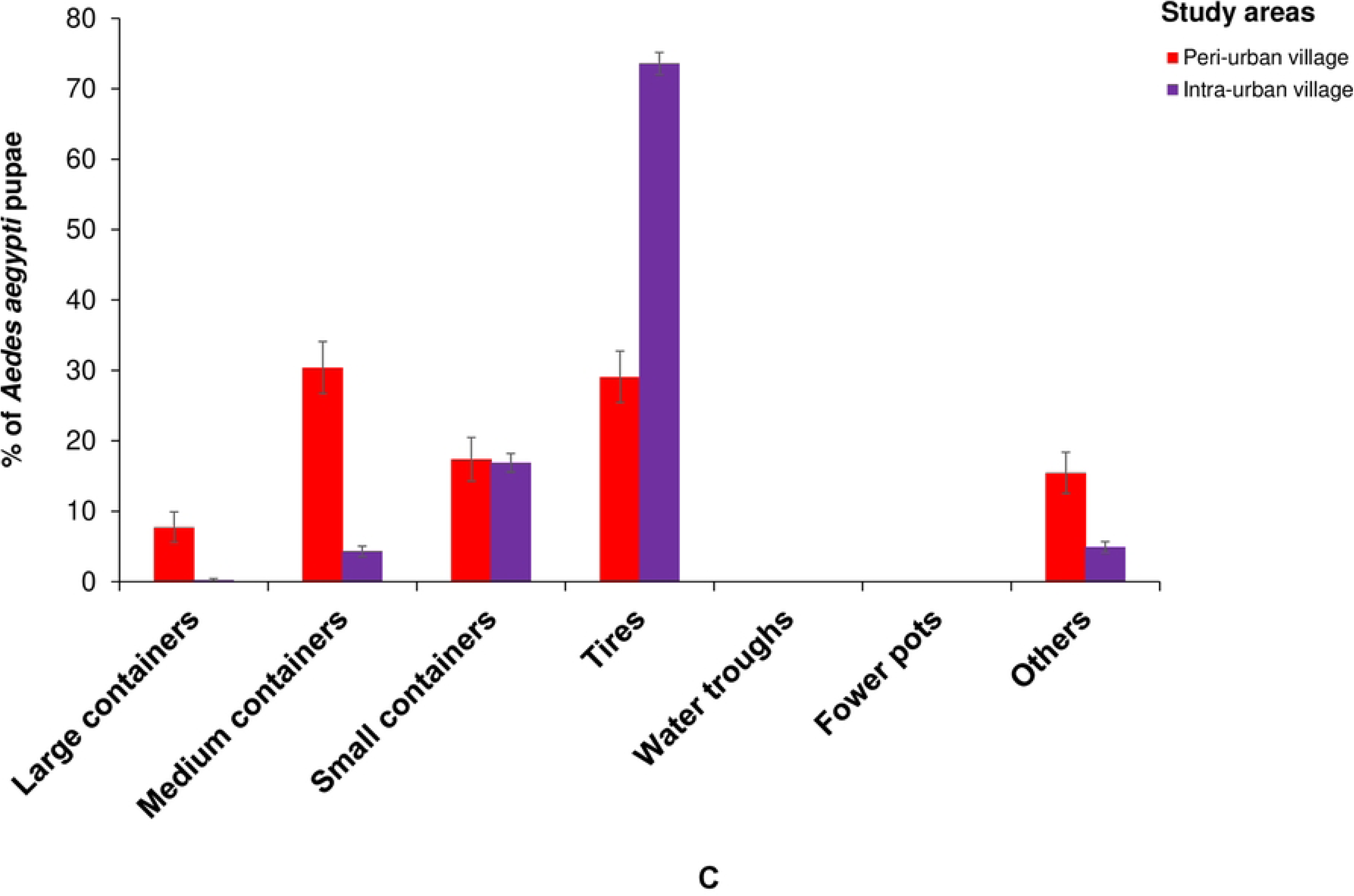
*Aedes aegypti* container productivity in the peri-urban and intra-urban villages of Cocody-Bingerville, southeastern Côte d’Ivoire from August 2023 to July 2024. A: Overall, B: Domestic ecozone, C: Peridomestic ecozone. Error bars indicate confidence intervals (95% CI). Others includes breeding containers made with brick holes, shoes, tarpaulins, wooden boxes, mortar, sheet metal, leaf armpits snail shells, underground puddles and tree holes.

**Table 3.**
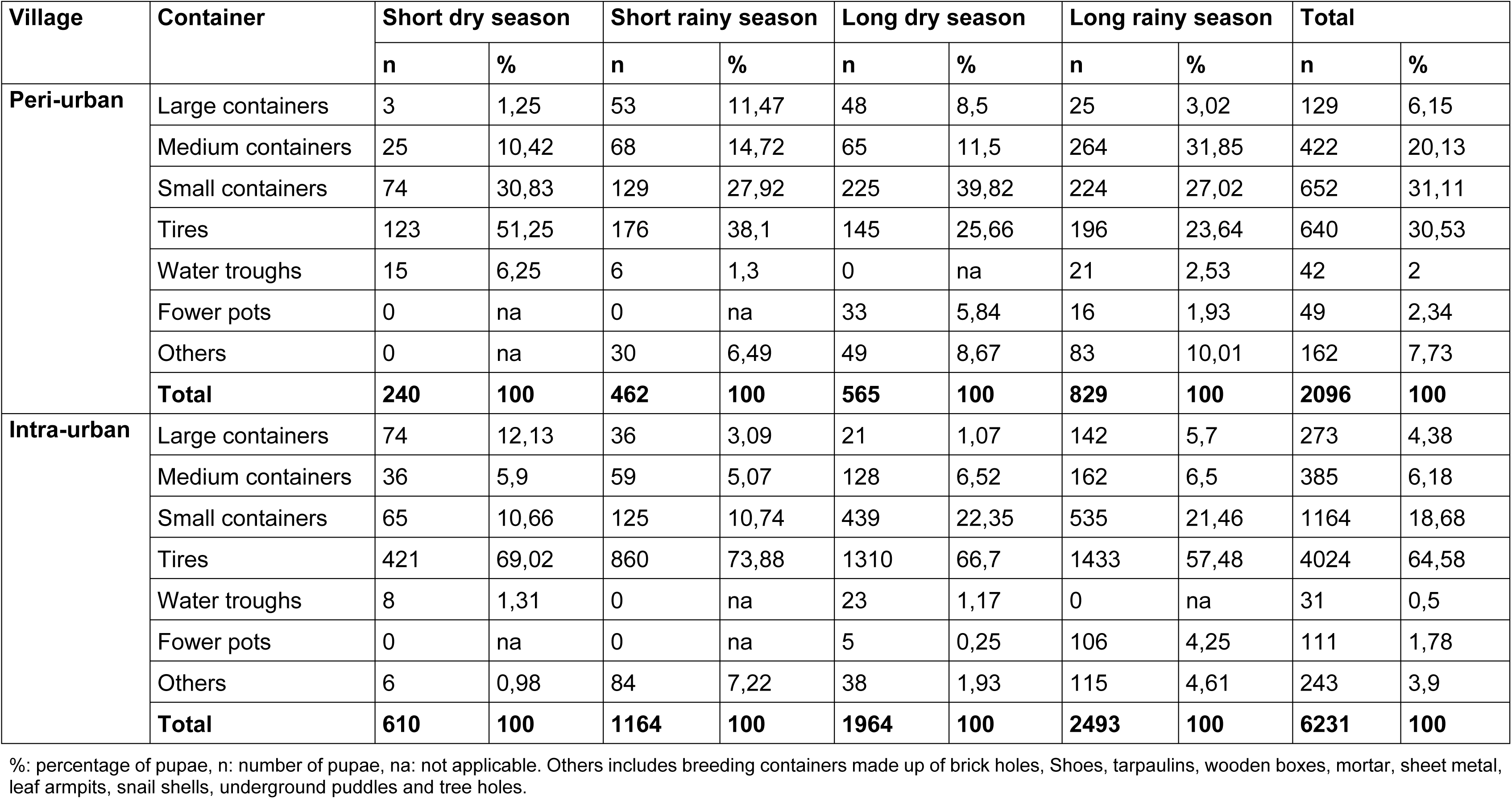
Seasonal variations in container productivity for Aedes aegypti pupae among peri-urban and intra-urban villages of Cocody-Bingerville, southeastern Côte d’Ivoire from August 2023 to July 2024.

Container productivity differed across the ecozones in all the study villages (χ^2^ = 742.89, df = 3, p < 0.0001). In the peri-urban villages, key containers were more productive in the domestic ecozones (70.9%, 1,487/2,096) compared with the peridomestic ecozones (29.1%, 609/2,096) (χ^2^ = 733.9, df = 1, p < 0.0001). The local most productive containers were small containers (36,72%, 546/1,487) followed by tires (31.14%, 463/1,487) and medium containers (15.94%, 237/1,487) in the domestic ecozones, and medium containers (30,38%, 185/609) followed by tires (29.06%, 177/609) and small containers in the peridomestic ecozones (Fig 6B and Fig 6C). In the intra-urban villages, key containers produced nearly similar proportions of pupae among the domestic ecozones (48.8%, 3,040/6,231) and peridomestic ecozones (51.2%, 3,191/6,231) (χ^2^ = 7.222, df = 1, p < 0.0001). Key containers were tires (55.10%, 1675/3,040) and small containers (73.61%, 2349/3,191) in the domestic ecozones, and tires (20.56%, 625/3,040) and small containers (16.89%, 539/3,191) in the peridomestic ecozones (Fig 6B and Fig 6C).

Container productivity was significantly influenced by seasonal variability in all the study villages (χ^2^ = 2247.5, df = 7, p < 0.0001). The highest numbers of pupae were recorded in LRS in all the study villages. During this period, the container productivity weas 3-fold lower in the peri-urban villages (829 pupae) than the intra-urban villages (2,493 pupae). The lowest numbers of pupae were found during SDS in the peri-urban villages (240 pupae) and the intra-urban villages (610 pupae). During LRS, the most productive containers were medium containers (31.85%, 264/829), small containers (27.02%, 224/829) and tires (23.64%, 196/829) in the peri-urban villages, and tires (57.48%, 1,433/2,493) and small containers (21.46%, 535/2,493) in the intra-urban villages. During this period, all key containers produced 82.51% (684/829) and 78.94% (1,968/2,493) of the pupae in the peri-urban intra-urban villages, respectively. During SDS, the highest pupal productivity was found in tires (51.25%, 123/240), small containers (30.83%, 74/240) and medium containers (10.42%, 25/240) in the peri-urban villages, and tires (69.02%, 421/610), large containers (12.13%, 74/610) and small containers (10.66%, 65/610) in the intra-urban villages (Table 3). During this period, key containers yielded 92.50% (222/240) in the peri-urban villages and (91.80%, 560/610) in the intra-urban villages.

### Aedes aegypti indices

#### Oviposition indices

OPI, MEO and EDI values were 39.95% [37.08%, 42.88%], 4.64 [4.08, 5.19] egg/ovitrap/week and 11.61 [10.49, 12.70] egg/ovitrap/week in the peri-urban villages, and (53.04% [50.04%, 56.04%]), 8.67 [7.89, 9.46] egg/ovitrap/week and 16.35 [15.19, 17.51] egg/ovitrap/week) in the peri-urban villages, respectively (S4 Table). GLMM showed that all the *Ae. aegypti* oviposition indices were statistically lower in the peri-urban villages than in the intra-urban villages (OPI: Estimate = 0.2835, Z = 44.88, p < 0.0001; MEO: Estimate = 0.6260, Z = 36.16, p < 0.0001; and EDI: Estimate = 0.3425, Z = 19.79, p < 0.0001). All oviposition indices were higher in the domestic ecozones than in the peridomestic ecozones in the peri-urban villages (all p < 0.05), but similar between the two ecozones in the intra-urban villages (all p > 0.05) (S5 Table). The three indices inconsistently varied over the seasons. OPI, MEO and EDI peaked in LDS, LDS and SRS in the peri-urban villages, and LDS, LRS and LRS in the intra-urban villages, respectively. OPI, MEO and EDI displayed the lowest values, respectively, during SRS, SDS and SDS in the peri-urban villages, and SRS, SRS and LDS in the intra-urban villages (S4 Table).

#### Stegomyia indices and epidemic risks of dengue and yellow fever

Fig 7 displays variations in the values of *Stegomyia* indices among the peri-urban and intra-urban villages and over the seasons. CI values were 29.96% [27.88%, 32.39%] in the peri-urban villages and 36.71% [34.50%, 38.96%] in the intra-urban villages (Fig 7A). HI values were estimated at 35.92% [32.63%, 38.11%] and 48.00% [45.14%, 50.87%] in the peri-urban and intra-urban villages, respectively (Fig 7B). BI values were 41.42 [38.04, 43.68] in the peri-urban villages and 56.17 [53.31, 59.00] in the intra-urban villages (Fig 7C). All *Stegomyia* indices were statistically comparable among the peri-urban and intra-urban villages (CI: F = 2.95, df = 1, p = 0.0999; HI: F = 3.072, df = 1, p = 0.0936; and BI: F = 2.602, df = 1, p = 0.121). *Stegomyia* indices varied significantly across the seasons in all the focus villages (all p < 0.001). CI displayed the highest values during LRS in both peri-urban (33.86% [29.76%, 38.14%]) and intra-urban (42.56% [38.52%, 46.69%]) villages. The lowest values of CI were observed during SDS (25.82% [20.75%, 31.42%]) in the peri-urban villages and during LDS (30.62% [24.41%, 35.08%]) in the intra-urban villages. HI showed peaks during LRS (51.00% [45.19%, 56.79%]) in the peri-urban villages and LRS (69.00% [63.43%, 74.19%]) in the intra-urban villages, that were statistically comparable (F = 18, df = 1, p = 0.5580). HI diminished gradually and exhibited lower values during SDS in the peri-urban (23.33% [18.66%, 28.54%]) and intra-urban (26.33% [21.44%, 31.70%]) villages. BI values peaked during LRS in the peri-urban villages (57.67 [51.67, 63.32]) and the intra-urban villages (83.00 [78.28, 87.07]). BI displayed the lower values during SDS in both peri-urban villages (23.67 [18.97, 28.89]) and intra-urban villages (29.67 [24.55, 35.19]).

**Fig 7.**
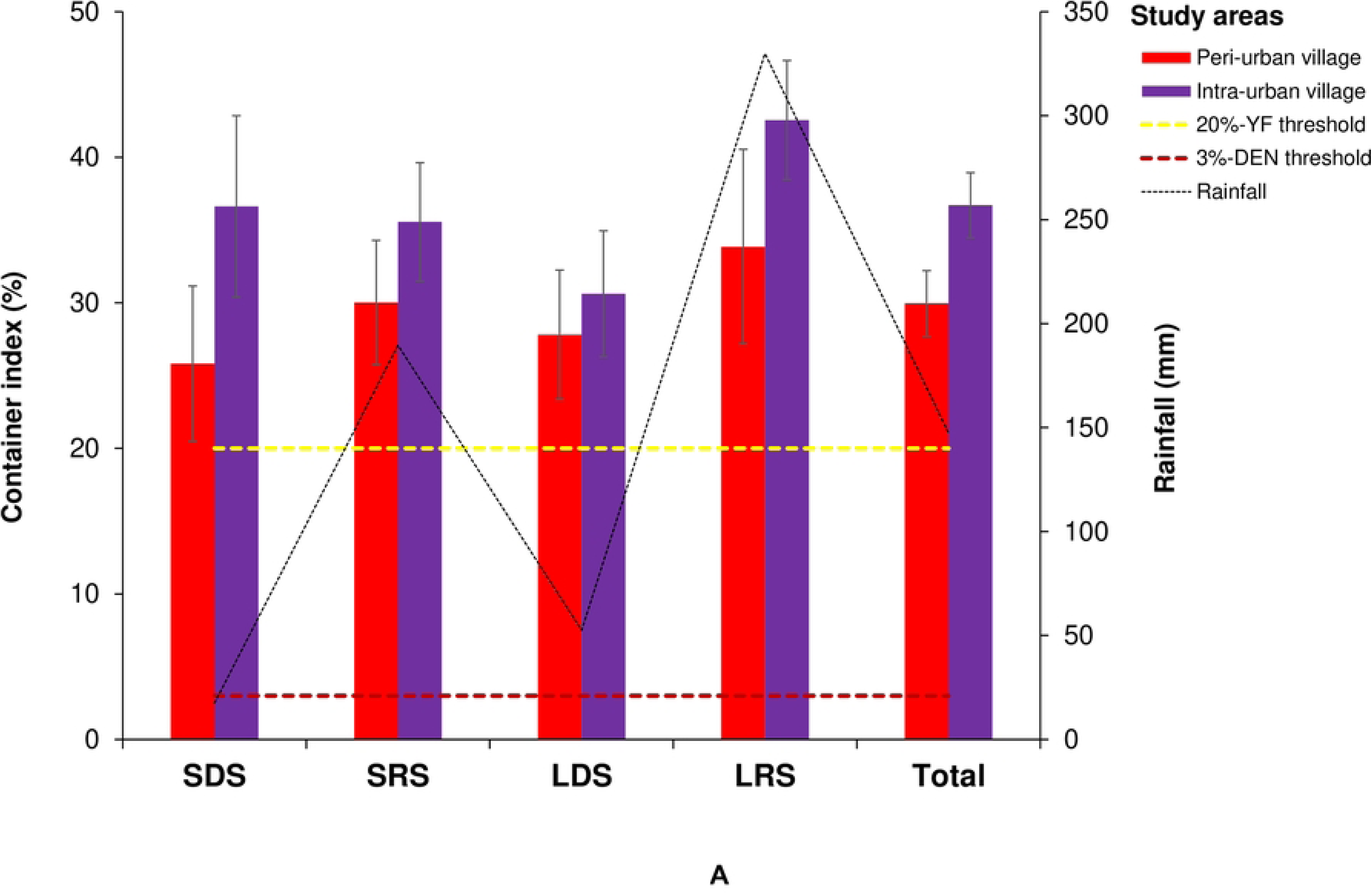
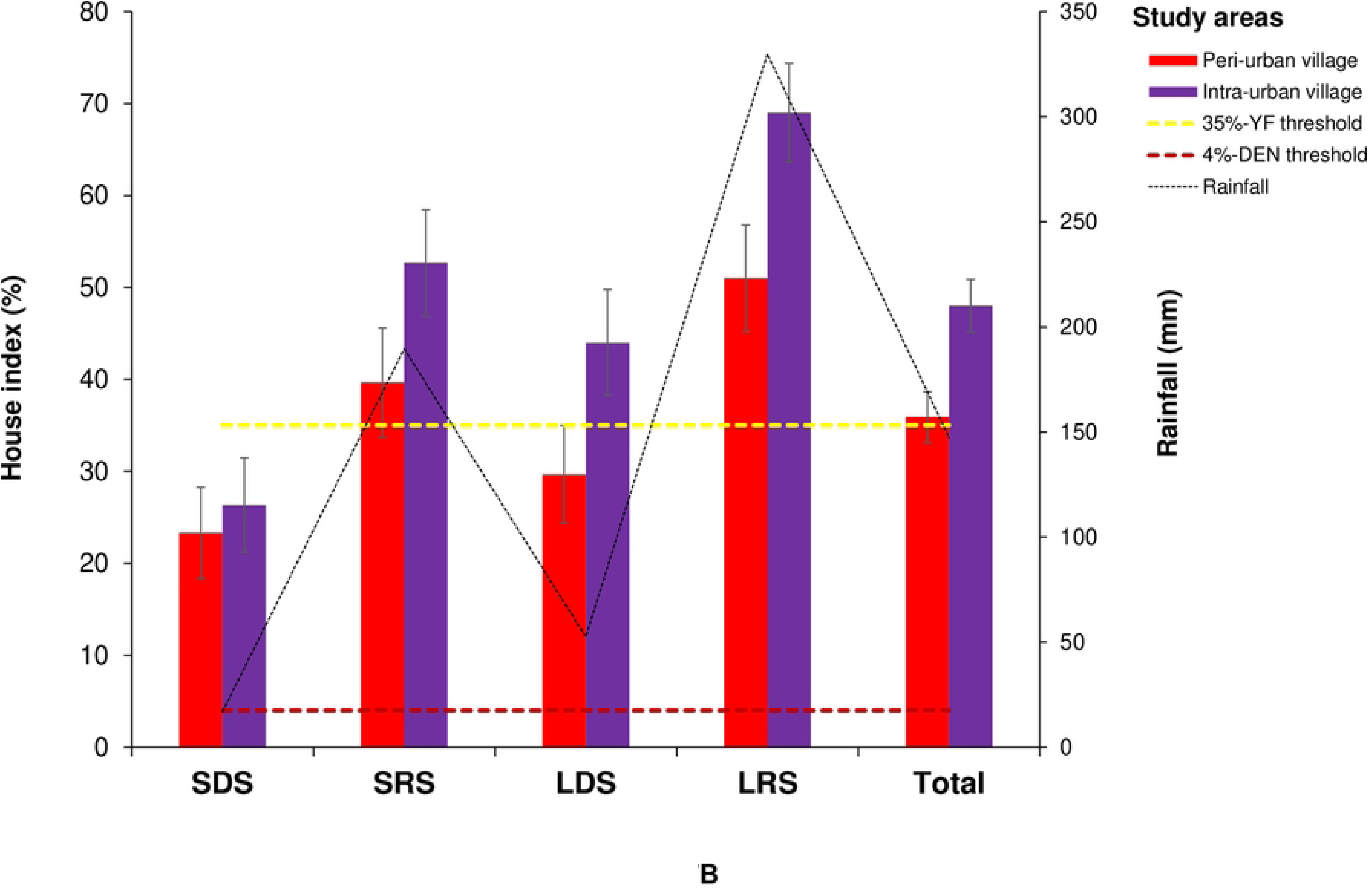
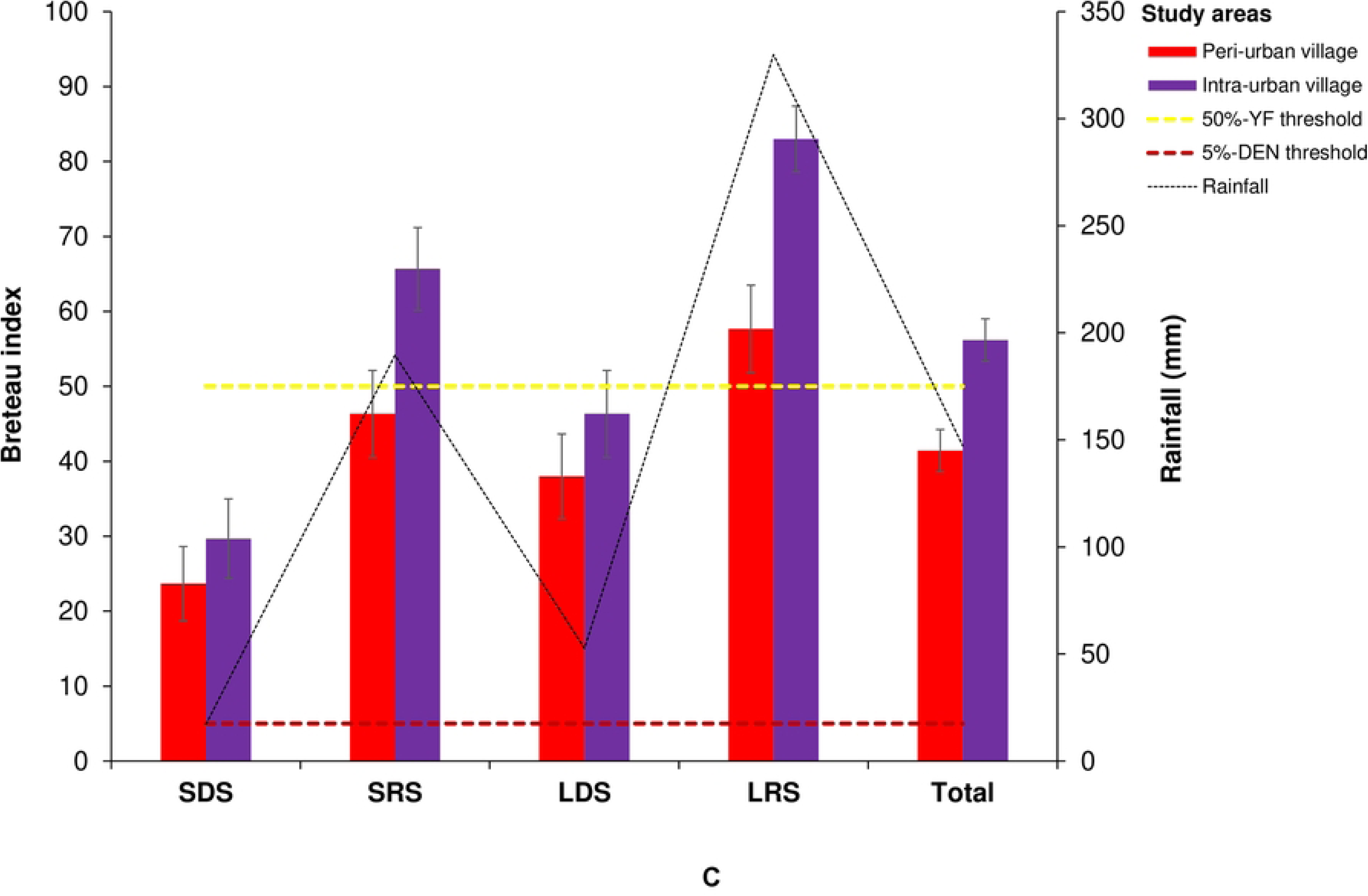
Seasonal variations of *Stegomyia* indices and epidemic risks of dengue and yellow fever in the peri-urban and intra-urban villages of Cocody-Bingerville, southeastern Côte d’Ivoire, from August 2023 to July 2024. %: percentage. Error bars show confidence intervals (95% CI). A: Container index, B: House index, C: Breteau index. LDS: long dry season, LRS: long rainy season, SDS: short dry season, SRS: short rainy season. Dengue epidemic thresholds are 4% for house index, 3% for container index and 5 for Breteau index [30]. Yellow fever epidemic thresholds are 35% for house index, 20% for container index and 50 for Breteau index [31].

Table 4 shows the level of the risks of DEN and YF epidemics associated with the *Stegomyia* indices recorded in the peri-urban and intra-urban villages. Overall, all CI, HI and BI values in the peri-urban and intra-urban villages exceeded the WHO DEN and YF epidemic thresholds [30,31], except for BI for YF in the peri-urban villages (Table 4). The risk indices corresponded to intervals of 5-8 for the peri-urban villages and 6-8 for the intra-urban villages and revealed high DEN and YF epidemic risk levels in all the study villages. Although DEN epidemic risk levels exhibited seasonal variations in all the peri-urban and intra-urban villages, values were permanently high and above the WHO DEN epidemic thresholds in all the study villages during the whole study period. YF epidemic risks were generally high in all the study villages during almost all seasons, but moderate during LDS and SDS in the peri-urban villages and SDS in the intra-urban villages.

**Table 4.** *Stegomyia* indices and levels dengue and yellow fever epidemic risks in the peri-urban and intra-urban villages of Cocody-Bingerville, southeastern Côte d’Ivoire, from August 2023 to July 2024.

| Village | Season | Container |  | House |  | Container index (%) |  | House index (%) |  | Breteau index |  | WHO density scale | Risk level |  |
| --- | --- | --- | --- | --- | --- | --- | --- | --- | --- | --- | --- | --- | --- | --- |
|  |  | Wet | Positive | Inspected | Positive | Mean | 95% CI | Mean | 95% CI | Mean | 95% CI |  | DEN | YF |
| Peri-urban | SDS | 275 | 71 | 300 | 70 | 25.82 | [20.75-31.42] | 23.33 | [18.66-28.54] | 23.67 | [18.97-28.89] | 4-7 | High | Moderate |
|  | SRS | 463 | 139 | 300 | 119 | 30.02 | [26.24-35.14] | 39.67 | [31.84-43.08] | 46.33 | [38.30-49.82] | 5-8 | High | High |
|  | LDS | 410 | 114 | 300 | 89 | 27.80 | [23.52-32.41] | 29.67 | [24.55-35.19] | 38.00 | [32.48-43.76] | 5-7 | High | Moderate |
|  | LRS | 511 | 173 | 300 | 153 | 33.86 | [29.76-38.14] | 51.00 | [45.19-56.79] | 57.67 | [51.67-63.32] | 6-8 | High | High |
|  | **Total** | **1659** | **497** | **1200** | **431** | **29.96** | **[27.88-32.39]** | **35.92** | **[32.63-38.11]** | **41.42** | **[38.04-43.68]** | **5-8** | **High** | **High** |
| Intra-urban | SDS | 243 | 89 | 300 | 79 | 36.63 | [30.55-38.96] | 26.33 | [21.44-31.70] | 29.67 | [24.55-35.19] | 4-8 | High | Moderate |
|  | SRS | 554 | 197 | 300 | 158 | 35.56 | [31.57-39.70] | 52.67 | [46.85-58.43] | 65.67 | [59.99-71.03] | 6-8 | High | High |
|  | LDS | 454 | 139 | 300 | 132 | 30.62 | [24.41-35.08] | 44.00 | [38.30-49.82] | 46.33 | [40.58-52.16] | 5-8 | High | High |
|  | LRS | 585 | 249 | 300 | 207 | 42.56 | [38.52-46.69] | 69.00 | [63.43-74.19] | 83.00 | [78.26-87.07] | 5-9 | High | High |
|  | **Total** | **1836** | **674** | **1200** | **576** | **36.71** | **[34.50-38.96]** | **48.00** | **[45.14-50.87]** | **56.17** | **[53.31-59.00]** | **6-8** | **High** | **High** |

%, percentage, DEN: dengue, YF: yellow fever, CI: confidence interval. LDS: long dry season, LRS: long rainy season, SDS: short dry season, SRS: short rainy season. *Stegomyia* indices are based on container infestation with *Aedes aegypti* larvae or pupae. WHO: World Health Organization. The WHO density scale and DEN and YF epidemic risk level are estimated according to the WHO guidelines [30,31].

#### Pupal indices

Table 5 shows the values of the pupal indices measured in the peri-urban and intra-urban villages. PCI and PHI respective values were significantly lower in the peri-urban villages (1.26 [1.02, 1.51] pupae/container and 1.75 [1.45, 2.05] pupae/house) compared with the intra-urban villages (3.39 [3.32, 3.46] pupae/container and 5.18 [4.44, 5.93] pupae/house) (PCI: F = 5.8535, df = 1, p < 0.0001; and PHI: F = 5.90, df = 1, p < 0.0001). Conversely, the respective values of PPI and HBI were statistically similar between the peri-urban villages (0.54 [0.52, 0.56] pupae/person and 0.27 [0.25, 0.28] female/person) and the intra-urban (1.25 [1.11, 1.30] pupae/person and 0.58 [0.56, 0.59] female/person) (PPI: F = 4.1279, df = 1, p = 0.0626; and HBI: F = 3.7898, df = 1, p = 0.0734).

**Table 5.**
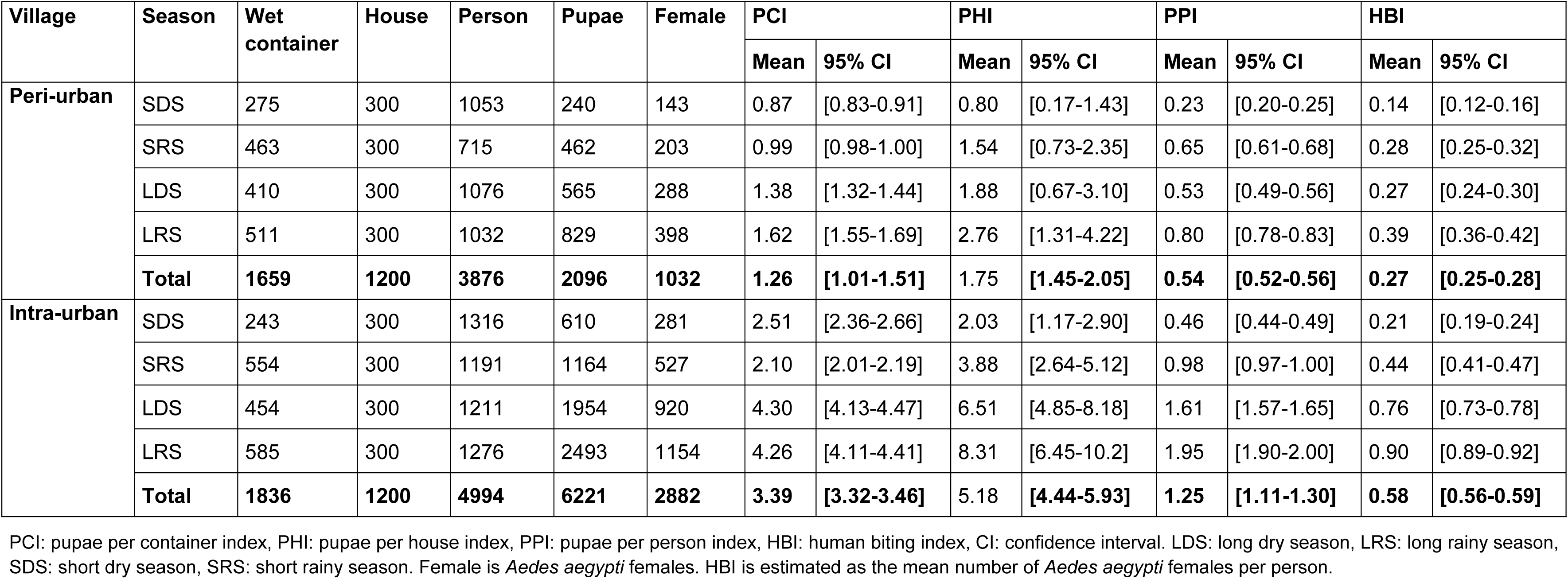
*Aedes aegypti* pupal indices in the peri-urban and intra-urban villages of Cocody-Bingerville, southeastern Côte d’Ivoire, from August 2023 to July 2024.

All the pupal indices followed the seasonal variations in all the study villages. PCI peaked during LRS in the peri-urban villages (1.62 [1.55, 1.69] pupae/container) and LDS in the intra-urban villages (4.30 [4.13, 4.47] pupae/container). PCI reached the lowest values during SDS (0.87 [0.83, 0.91] pupae/container) in the peri-urban and SRS (2.10 [2.01, 2.19] pupae/container) in the intra-urban villages. PHI exhibited peaks during LRS in both peri-urban (2.76 [1.31, 4.22] pupae/house) and intra-urban (8.31 [6.45, 10.20] pupae/house) villages, and reached the lowest values during SDS in the peri-urban villages (0.80 [0.17, 1.43] pupae/house) and intra-urban villages (2.03 [1.17, 2.90] pupae/house). PPI peaks occurred simultaneously during LRS in both peri-urban villages (0.80 [0.78, 0.83] pupae/person) and intra-urban villages (1.95 [1.90, 2.00] pupae/person). PPI diminished considerably across seasonal variations and attained the lowest values during SDS in the peri-urban villages (0.23 [0.20, 0.25] pupae/person) and the intra-urban villages (0.46 [0.44, 0.49] pupae/person). HBI peaked during LRS in the peri-urban villages (0.39 [0.36, 0.42] female/person) and the intra-urban villages (0.90 [0.89, 0.92] female/person). HBI decreased over the seasons and showed the lowest values during SDS in both peri-urban villages (0.14 [0.12, 0.16] female/person) and in intra-urban villages (0.21 [0.19, 0.24] female/person).

### Climate effects on *Aedes aegypti* abundance and arbovirus transmission risks

We found weak, but positive and statistically significant correlations between *Ae. aegypti* numbers and rainfall in the peri-urban villages (ρ = 0.1913, p < 0.0001) and the intra-urban villages (ρ = 0.068, p < 0.0001). *Aedes aegypti* numbers and temperature exhibited negative and weak associations that were significant in the peri-urban villages (ρ = - 0.1474, p < 0.0001), but not significant in the intra-urban villages (ρ = - 0.0219, p = 0.4394). *Aedes aegypti* numbers and RH showed weak and non-significant correlations which were positive in the peri-urban villages (ρ = 0.008, p = 0.8052) and negative in the intra-urban villages (ρ = -0.0053, p = 0.8528).

Among *Stegomyia* indices, HI and BI were positively, strongly and significantly correlated with rainfall in the peri-urban and intra-urban villages (all p < 0.05) (S6 Table). For pupal indices, PHI, PPI and HBI were positively, moderately and significantly correlated with rainfall in the peri-urban villages (all p < 0.05). However, the corrections between rainfall and all pupal indices were positive, weak and non-significant in the intra-urban villages (all p > 0.05) (S6 Table). The temperature was negatively, weakly and non-significantly associated with all *Stegomyia* indices and all pupal indices in all the study villages (all p > 0.05), except for HI in the peri-urban villages and HI and BI in the intra-urban villages (S6 Table). Overall, there were negative, weak and non-significant relationships between RH and all *Stegomyia* indices and all pupal indices in the peri-urban and intra-urban villages (all p > 0.05) (S6 Table).

## Discussion

This study provides a comparative analysis of *Ae. aegypti* populations and epidemic risk levels in the peri-urban and intra-urban villages of Cocody-Bingerville, an area accounting for the large majority (80-90%) of DEN and YF cases reported in Côte d’Ivoire [10,11,15,17]. While previous entomological investigations and outbreak responses have predominantly focused on urbanized neighborhoods, including the intra-urban villages [6,19], the peri-urban villages have remained understudied. Our present study showed that the peri-urban and intra-urban villages were heavily infested with *Ae. aegypti* immatures and productive containers, and both had similarly high *Stegomyia* indices that exceeded the WHO-established DEN and YF epidemic thresholds. This demonstrates that residents in the peri-urban and intra-urban villages face equally high risks of the outbreaks. Thus, the findings presented here highlight distinct spatial and seasonal patterns of *Ae. aegypti* immature ecologies, underlining the importance of integrating the peri-urban villages into vector surveillance and control strategies.

Our data showed that *Ae. aegypti* species was the most abundant *Aedes* species in both peri-urban villages (98.1%) and intra-urban villages (99.8%). The high dominance of *Ae. aegypti* in these villages could be explained by high density of human populations and high number of suitable artificial containers (e.g., tires, small containers, medium containers, large containers, flowerpots and water troughs). Indeed, *Ae. aegypti* is a highly anthropophagic mosquito that preferentially feeds on humans and breeds in man-made containers [21,33]. Moreover, the local climatic conditions (e.g., rainfall (149 mm), temperature (28 °C) and RH (83%) are likely favorable for *Ae. aegypti* oviposition and egg hatching, larval development and pupation [8,34]. However, *Ae. aegypti* proportions were significantly lower in the peri-urban than in the intra-urban villages. Indeed, both human population size and number of positive containers were lower in the peri-urban villages compared to the intra-urban villages. This could be potentially attributed to varying proximity of the study villages to the highly urbanized Cocody-Bingerville. Indeed, while the intra-urban villages lie within Cocody-Bingerville, the peri-urban villages are situated 5-15 km away.

We found that local containers were highly infested with *Ae. aegypti* immatures in both peri-urban and intra-urban villages. The most *Ae. aegypti*-productive containers were small containers, tires and medium containers in the peri-urban villages, and tires and small containers in the intra-urban villages. Taken together, these key containers produced 82% of the pupae in the peri-urban villages and 83% of the pupae in the intra-urban villages. Our results are consistent with those reported from urbanized areas in Côte d’Ivoire (e.g., Abidjan) and other African countries (e.g., Burkina Faso and Kenya) where artificial containers (e.g., tires, discarded cans, water storage receptacles) were found to harbor high numbers of *Ae. aegypti* larvae or pupae [18,35–37]. Oviposition indices were all significantly lower in the peri-urban villages than in the intra-urban villages. This difference could be probably due to a differential ovipositing adaptation in *Ae. aegypti* females and their proportions [33]. The massive presence of positive containers in the peri-urban and intra-urban villages could be probably due to the spillover effects of urbanization coupled with a mismanagement of water holding containers and solid waste and poor environmental sanitation and hygiene [6]. However, the numbers of containers were lower in the peri-urban villages compared with the intra-urban villages. Indeed, the peri-urban villages seem to be less influenced by urbanization than the peri-urban villages that are more distant from urban Cocody-Bingerville. *Aedes aegypti* abundance and container productivity showed ecozonal and seasonal shifts among the study villages. Indeed, *Ae. aegypti* proportions were higher in the domestic ecozones than in the peridomestic ecozones in the peri-urban villages, but comparable among the ecozones in the intra-urban villages. This disproportional distribution is probably likely to the differences in the distributional patterns of larval containers among the ecozones [6,19]. Indeed, the productive containers were more in the domestic ecozones than in the peridomestic ecozones in the peri-urban villages, but were equally distributed among the two ecozones in the intra-urban villages. The high presence of *Ae. aegypti* within and around houses could raise the risk of transmission of DEN and YF viruses to the inhabitants in both study villages [6,19,38]. Local *Ae. aegypti* abundance shifted over seasonal variations, displaying higher proportions during LRS and lower proportions during SDS in both peri-urban and intra-urban villages. Similarly, key container productivity was, respectively, 83% and 79% in the peri-urban and intra-urban villages during the most container productivity period (i.e., LRS), and 93% and 92% in the peri-urban and intra-urban villages during the least container productivity period (i.e., SDS). As the local temperature (23.5-34.9 °C) and RH (64.7-84.9%) varied little across the seasonal variability and ranged within optimal conditions for *Ae. aegypti*, rainfall appears to be the main driver of the local population dynamics. High rainfall recorded during LRS (329.9 mm) may have flooded many local containers providing high number of aquatic oviposition grounds to gravid females, thus resulting in the proliferation of *Ae. aegypti* populations. Conversely, the lower numbers of *Ae. aegypti* during SDS (17.05 mm) might be due to a decline of rainfall associated with a dry out of containers in the study villages [39]. Surprisingly, *Ae. aegypti* abundance was significantly higher during LDS compared with SDS and SRS in the peri-urban and intra-urban villages. This could be explained by unexpected meteorological variations due to climate change as rainfall was relatively high during LDS (52.55 mm). Moreover, people might probably store water for long duration due to drought that could increase provably container productivity [6,19].

Local community-led larval source management programs targeting the identified key containers in appropriate ecozones during SDS could be effective for preventing an increase in *Ae. aegypti* numbers during LRS. Such targeted interventions could help contain current and prevent future outbreaks of DEN and YF for lower human, logistical and financial resources.

All the *Stegomyia* indices exceeded the WHO DEN and YF epidemic thresholds, and corresponded to the WHO risk intervals of 5–8 in the peri-urban villages and 6–8 in the intra-urban villages. The *Stegomyia* indices were statistically similar among the peri-urban and intra-urban villages. In particular, the high-risk levels observed in the peri-urban villages were further compounded by the presence of five additional sylvatic *Aedes* species, including *Ae. vittatus* that is a competent vector for both DENV and YFV. These data underscores the elevated epidemic risks in all the study villages, which is a concern often seen in major African urban centers (e.g., Abidjan, Ouagadougou, and Nairobi) [19,35,40], but rarely in peri-urban villages. Therefore, the high risks of DEN and YF outbreaks found in the peri-urban and intra-urban villages are exceptional and might require particular attention. This could be mediated by the driving effects of the rapid and unplanned urbanization of Cocody-Bingerville. For instance, increased availability of humans and containers associated with warm and humid climate could offer ideal conditions for the proliferation of *Ae. aegypti* populations in these villages. Worryingly, *Stegomyia* indices varied slightly across seasons, with DEN epidemic risks remaining permanently high and above the WHO epidemic thresholds year-round in both peri-urban and intra-urban villages. YF epidemic risks were generally high across most seasons, except for moderate risks observed during LDS and SDS in the peri-urban, and during SDS in the intra-urban villages. These seasonal variations emphasize the importance of year-round vector surveillance and control measures, as the risks do not significantly decrease during less productive periods. This could explain ongoing and recurrent DEN outbreaks, often coupled with sporadic YF cases, in Cocody Bingerville.

As pupal mortality is low [41], pupal indices could thus reflect *Ae. aegypti* adult density and female biting patterns. In the present study, PCI and PHI were significantly lower in the peri-urban villages than in the intra-urban villages, whereas PPI and HBI were statistically equal between all the focus villages. Local *Ae. aegypti* populations may have strong anthropophilicity and resize pupal productivity and emerging biting females to the number of available human hosts. The high abundance of pupae in the study villages suggests that local biotic (i.e., human host) and abiotic (i.e., rainfall, temperature and RH) factors appear suitable for larval development, pupation and adult emergence [42]. Emerged females are expected to bite and transmit DENV and YFV to the local people. This indicates similar potential for human-vector contacts, thereby maintaining consistent levels of DEN and YF transmission risk across both study settings.

Complementary investigations are required to address some limitations and gain a better understanding of arboviral outbreaks. Analyzing DENV and YFV in *Aedes* and human populations is important to detect their active circulation. In parallel, serological studies in people are needed to assess prior exposure and current transmission dynamics through the detection of specific antibodies. Characterizing biological interactions (predation), food inputs and physicochemical parameters of water-container systems should help to better understand container infestation and productivity. Knowing *Ae. aegypti* adult females’ biting behaviors (indoors *vs.* outdoors) and period (daytime *vs.* nighttime), resting places (indoors *vs.* outdoors) and host preferences (humans *vs.* animals) is crucial to design effective vector control plans. Finally, understanding local community knowledge and practices regarding vector control and solid waste management will be critical for the successful implementation of sustainable, community-based control programs.

Above all, the present study, conducted during an ongoing DEN outbreak in Cocody-Bingerville, provides valuable and timely insights into the eco-epidemiology of *Ae. aegypti* in both peri-urban and intra-urban villages. The ongoing outbreak offers a unique opportunity to assess vector ecology and arbovirus transmission dynamics, under real-world conditions, allowing for more detailed and context-specific understanding of local *Ae. aegypti* populations and their role in disease epidemics. Our findings indicate that both peri-urban and intra-urban villages are exposed to consistently high epidemic risks, with *Stegomyia* indices surpassing the WHO-established thresholds across all seasons. This is particularly significant given that such high-risk levels are typically associated with more urbanized areas. This highlights the need to expand *Aedes* vector surveillance and control to peri-urban villages, which have historically received less attention despite the potential for these areas to act as reservoirs for transmission that can reintroduce the virus into more densely populated urban centers. The current study introduced, for the first time in Côte d’Ivoire, OPI, MEO and EDI indicators to evaluate *Ae. aegypti* oviposition activities. This could enhance arboviral risk assessment as ovitrap-based surveillance is simpler in use, less resource-demanding, less time-consuming and cheaper than larval and pupal surveys. The peri-urban villages should be also considered when developing and implementing arboviral outbreak responses as these neglected settings could be a source of failure of *Aedes* vector control interventions operating in cities. This study underscores the importance of integrating vector control efforts that are specifically adapted to the local context, particularly in peri-urban villages that are undergoing rapid, often unplanned, urbanization. The presence of high densities of productive containers, coupled with favorable climatic conditions, creates an ideal environment for *Ae. aegypti* populations to thrive, thereby amplifying the risk of DEN and YF outbreaks. As shown in the urbanized cities of Ouagadougou and Abidjan [43–45], integrated community-led larval source management interventions targeting the incriminated key containers (i.e., tires, small containers and medium containers) in appropriate ecozones (i.e., domestic and/or peridomestic environment) and during identified less production season (i.e., SDS) could be cost-effective for containing the current DEN outbreaks and preventing future arboviral epidemics in the peri-urban and intra-urban villages of Cocody-Bingerville.

## Conclusion

Given the context of the ongoing DEN outbreak in Cocody-Bingerville, the study provided real-time data on vector dynamics and the seasonality of container productivity. Indeed, both peri-urban and intra-urban villages were found mostly infested with *Ae. aegypti* immatures. Over 80% of pupae was produced by a few types of aquatic habitats (i.e., tires, small containers and medium containers). Containers’ productivity was higher in the domestic ecozones in the peri-urban villages, but similar among the domestic and peridomestic ecozones in the intra-urban villages, was lowest during SDS in all the study villages. DEN and YF epidemic risks exceeded the WHO thresholds throughout. These insights are especially important for informing vector control strategies during active outbreaks. We will further analyze *Aedes* samples for DENV and YFV to detect their active circulation. Taken together, the results of this study suggest that the peri-urban villages should be prioritized in the national arboviral outbreak responses. Integrated community-based larval source management programs, particularly targeting key containers during the identified low-productive season in high-productive ecozones, could be highly cost-effective in reducing *Aedes* densities and controlling the ongoing and preventing future outbreaks of DEN and YF in Cocody-Bingerville.

## Supporting information

**S1 Table. Species composition of mosquito adults emerged of eggs, larvae and pupae collected in the peri-urban and intra-urban villages of Cocody-Bingerville, southeastern Côte d’Ivoire, from August 2023 to July 2024.**

**S2 Table. Geographical distribution of *Aedes aegypti* in the peri-urban and intra-urban villages of Cocody-Bingerville, Côte d’Ivoire, from August 2023 to July 2024.**

**S3 Table. Seasonal variations in *Aedes aegypti*-positive breeding sites in the peri-urban and intra-urban villages of Cocody-Bingerville, southeastern Côte d’Ivoire, from August 2023 to July 2024.**

**S4 Table. *Aedes aegypti* oviposition indices in the peri-urban and intra-urban villages of Cocody-Bingerville, southeastern Côte d’Ivoire, from August 2023 to July 2024.**

**S5 Table. *Aedes aegypti* oviposition indices across ecozones in the peri-urban and intra-urban villages of Cocody-Bingerville, southeastern Côte d’Ivoire, from August 2023 to July 2024.**

**S6 Table. Correlations between *Stegomyia* indices, pupal indices and climate variables in the peri-urban and intra-urban villages of Cocody-Bingerville, southeastern Côte d’Ivoire, from August 2023 to July 2024.**

## Acknowledgements

The authors thank the administrative and traditional authorities who authorized this research activities and the inhabitants of all the study areas.

## Author contributions

**Conceptualization:** JZBZ, GDM, LSD, JFM, AAA, SB, SCB.

**Data curation:** YNB, JZBZ, JDKD, PNC.

**Formal analysis:** YNB, JZBZ.

**Funding acquisition:** JZBZ, GDM, LSD, JFM, AAA, SB, SCB.

**Investigation:** YNB, JZBZ, JDKD, MT.

**Methodology:** YNB, JZBZ, FH, LSD, JFM, SCB, MT.

**Project administration:** JZBZ, FH, SB, SCB.

**Data collection:** YNB, JDKD, PNC.

**Resources:** JZBZ, MT.

**Supervision:** JZBZ, MT.

**Visualization:** YNB, JZBZ, JDKD.

**Writing – original draft:** YNB, JZBZ.

**Writing – review & Editing:** YNB, JZBZ, PNC, PGM, FH, GDM, LSD, JFM, AAA, SB, SCB, MT.

